# *TGIF1* is required for chicken ovarian cortical development and generation of the juxtacortical medulla

**DOI:** 10.1101/2021.03.30.437645

**Authors:** Martin Andres Estermann, Claire Elizabeth Hirst, Andrew Thomas Major, Craig Allen Smith

## Abstract

During early embryogenesis in amniotic vertebrates, the gonads differentiate into either ovaries or testes. The first cell lineage to differentiate gives rise to the supporting cells; Sertoli cells in males and pre-granulosa cells in females. These key cell types direct the differentiation of the other cell types in the gonad, including steroidogenic cells. The gonadal surface epithelium and the interstitial cell populations are less well studied, and little is known about their sexual differentiation programs. Here, we show the requirement of the transcription factor gene *TGIF1* for ovarian development in the chicken embryo. *TGIF1* is expressed in the two principal ovarian somatic cell populations, the cortex and the pre-granulosa cells of the medulla. *TGIF1* expression is associated with an ovarian phenotype in sex reversal experiments. In addition, targeted over-expression and gene knockdown experiments indicate that TGIF1 is required for proper ovarian cortical formation. *TGIF1* is identified as the first known regulator of juxtacortical medulla formation. These findings provide new insights into chicken ovarian differentiation and development, specifically in the process of cortical and juxtacortical medulla formation, a poorly understood area.

**SUMMARY STATEMENT:** The transcription factor TGIF1 is required for proper ovarian sex differentiation in chicken embryos, regulating development of the cortical and juxtacortical medulla, independently of the supporting cell sex lineage.

## INTRODUCTION

Vertebrate gonadal sex differentiation is a unique process whereby the embryonic gonadal primordium typically adopts either an ovarian or testicular fate (Brennan and Capel, 2004, Stevant and Nef, 2019, Rotgers et al., 2018). This process involves the expression of sexually dimorphic genes that activates one pathway and represses the other, making testis and ovary formation mutually exclusive (Kim et al., 2006, Li et al., 2017). The undifferentiated gonad initially comprises the same set of uncommitted cell lineage precursors; so-called supporting cells, steroidogenic progenitors, germ cells and some other less well-defined cells (Lin et al., 2017, Lin and Capel, 2015, Nef et al., 2019). The first cell lineage to differentiate is the supporting cell lineage, giving rise to Sertoli cells in the male gonad and pre-granulosa cells in females (Niu and Spradling, 2020, Chen et al., 2017, Zhang et al., 2015a). These cells are then thought to direct other lineages down the testicular or ovarian pathways, respectively (Lin and Capel, 2015, Rotgers et al., 2018, Wear et al., 2017, Gustin et al., 2016). In males, Sertoli cells organize into testis cords and signal to neighboring steroidogenic precursors to become sex steroid-hormone producing fetal Leydig cells in the developing testis (Yao et al., 2002). The same lineage gives rise to thecal cells in the developing ovary, although this requires interactions with the germ cells (Liu et al., 2015, Stevant et al., 2019). Germ cells themselves follow a fate governed by signals from the somatic component of the gonad, giving rise to spermatogonia in the testis and oogonia in the ovary (Barrios et al., 2010, Bowles et al., 2010, DiNapoli et al., 2006, Spiller et al., 2017). Other cell types in the embryonic gonad are less well characterized, including the gonadal surface epithelium (the source of the supporting and some of the steroidogenic cell lineages in mouse) and non-steroidogenic “interstitial” cells derived from the surface epithelium or the adjacent mesonephric kidney (DeFalco et al., 2011, Rotgers et al., 2018, Svingen and Koopman, 2013, Stevant and Nef, 2019).

Gonadal sex differentiation has been widely studied as a paradigm for the molecular genetic regulation of development. In the mouse model, Y chromosome linked *Sry* gene initiates the testis developmental program (Koopman et al., 1991, Sinclair et al., 1990, Hacker et al., 1995, Kashimada and Koopman, 2010). It activates the related *Sox9* gene, leading to Sertoli cell differentiation, and subsequent downstream singling to channel other cell types down the male pathway (Sekido et al., 2004, Sekido and Lovell-Badge, 2008, Qin and Bishop, 2005, Li et al., 2014, Gonen et al., 2017). In mouse, once Sry has activated Sox9, the latter can drive complete testis formation (Qin and Bishop, 2005), through activation of Fgf9 signaling and other mechanisms (Vidal et al., 2001, Kim et al., 2007, Gonen and Lovell-Badge, 2019, Schmahl et al., 2004, Colvin et al., 2001). In female mammals (genetically XX), the absence of *Sry* allows activation of the signaling molecule R-Spondin1, Wnt4, and stabilization of β-catenin, and downstream expression of the transcription factor, Foxl2 (Li et al., 2017, Parma et al., 2006, Tomizuka et al., 2008, Maatouk et al., 2008, Chassot et al., 2008, Jordan et al., 2003). This engages the ovarian pathway. Genetic antagonism exists throughout these opposing testis and ovarian pathways; Fgf9 (male) vs Wnt4 (female), for example, and Sry vs R-Spo1 (Kim et al., 2006, Lavery et al., 2012, Lau and Li, 2009). However, the molecular regulation of gonadal sex differentiation is still incompletely understood, specifically with regard to cell types other than the key supporting cell lineage. Recently, bulk and single-cell RNA sequencing approaches have expanded the list of genes implicated in gonadal sex differentiation (Stevant et al., 2019, Stevant et al., 2018, Estermann et al., 2020). Many novel genes uncovered by these approaches remain to be functionally analyzed.

Our understanding of vertebrate gonadal development has been enhanced through comparative studies in non-mammalian models. While several core genes required for gonadal sex differentiation are conserved across species (Sox9 in the testis and Foxl2 in the ovary, for example) (Kent et al., 1996, Major et al., 2019, Capel, 2017), upstream master sex genes can be divergent. Sry is absent on non-mammals, and so other master sex triggers must exist. Among egg-laying vertebrates, the transcription factor DMRT1 plays a major role, analogous to Sry. DMRT1 acts as a master sex switch in birds and in many reptiles with temperature dependent sex determination, inducing testis development (Smith et al., 2009, Ioannidis et al., 2020, Sun et al., 2017, Lambeth et al., 2014). The chicken embryo, in particular, has proved valuable insights in the genetic reregulation of gonadal sex differentiation, the evolution of genetic sex switches, and the cell biology of gonadogenesis (Sekido and Lovell-Badge, 2007, Guioli et al., 2020, Smith and Sinclair, 2004, Estermann et al., 2020). As embryonic development occurs *in ovo* and is accessible for experimental manipulation, the chicken provides a powerful model for functional analysis of gonadal sex-determining genes (Schmid et al., 2015). This model has been particularly useful for elucidating the cellular events underpinning gonad formation. Chickens have a ZZ male; ZW female sex chromosome system, in which Z-linked *DMRT1* gene operates as a master testis regulator via a dosage mechanism (two doses in males) (Ioannidis et al., 2020). In ZZ embryos, the gonads differentiate into bilateral testes. As in mammals, the seminiferous cords form in the inner gonadal medulla in chicken, comprising Sertoli that enclose germ cells (Smith and Sinclair, 2004). The male germ cells undergo mitotic arrest, entering meiosis only after hatching (Ayers et al., 2013). In the female chicken gonad, the inner medulla is the site of aromatase gene expression. Aromatase catalyzes the synthesis of estrogens, which are essential for ovarian differentiation in birds (and other egg laying vertebrates) (Scheib, 1983, Vaillant et al., 2001b, Pieau and Dorizzi, 2004).

The avian model is particularly useful for shedding light on the role of the gonadal surface epithelium. In mouse, the surface epithelium gives rise to the supporting cell lineage and then contributes to the steroidogenic lineage (Lin et al., 2017, Stevant et al., 2018, Stevant et al., 2019). In chicken, the surface epithelium gives rise to non-steroidogenic interstitial cells, not the supporting cell lineage as in mouse (Estermann et al., 2020, Sekido and Lovell-Badge, 2007). Prior to gonadal sex differentiation in chicken, the left gonadal epithelial layer is thicker than that the right one (in both sexes) (Omotehara et al., 2017, Guioli et al., 2014). During sex differentiation, this asymmetry becomes less marked in males (Guioli et al., 2014). However, symmetry is maintained and becomes very pronounced in females (Smith and Sinclair, 2004). The right gonad regresses in female birds, whereas the epithelium of the left gonad continues to proliferate to become a thickened cortex (Guioli et al., 2014). Increased proliferation in the left cortex, rather than increased apoptosis in the right cortex, is primarily responsible for the observed asymmetric cortical development (Ishimaru et al., 2008). The left cortex is critical to ovarian development in the avian model. The thickened left cortex contains both somatic cells and proliferating germ cells that enter meiosis to later arrest at prophase I (Ukeshima, 1996). Immediately beneath the cortex of the left ovary, interstitial medullary cells form a compact region called the juxtacortical medulla (JCM). We have previously shown through single-cell RNA-seq that the cells of the JCM are non-steroidogenic and derive from the ovarian surface epithelium (Estermann et al., 2020). The functional significance of the JCM is unclear, although at later stages it expresses enzymes involved in retinoic metabolism, and retinoic acid is implicated in cortical germ cell meiosis (Smith et al., 2008).

We previously conducted bulk RNA-sequencing to identify novel genes involved in development of the chicken ovary (Ayers et al., 2015). This screen identified *TGIF1* (*TGF-β Induced Factor Homeobox 1*). *TGIF1* encodes a homeobox transcription factor that belongs to the superfamily of TALE homeodomain proteins known to control many developmental processes, including gastrulation, cell proliferation, and differentiation (Wotton et al., 1999a, Wotton et al., 1999b, Lorda-Diez et al., 2009, Melhuish and Wotton, 2000, Wotton et al., 2001, Liu et al., 2014, Powers et al., 2010). It has not previously been associated with gonadal sex differentiation in any species. In the current study, we describe the role of *TGIF1* in chicken ovarian development. *TGIF1* is specifically upregulated in female gonads at the onset of sexual differentiation, expressed in cortical and pre-granulosa cells and is associated with the ovarian phenotype. Over-expression and knockdown of *TGIF1* show that it is required for the formation of the female cortex and the juxtacortical medulla. The data suggest that TGIF1 is required for proper ovarian development in the avian model, acting downstream of estrogen signaling.

## RESULTS

### *TGIF1* but not *TGIF2* shows sexually dimorphic expression in embryonic chicken gonads

TGIF1 was firstly identified as a candidate gene in avian gonadal sex differentiation from a gonadal RNA-seq performed in our laboratory (Ayers et al., 2015). Differential expression analysis showed that *TGIF1* mRNA expression was significantly higher in female compared to male gonads at the onset of sex differentiation (Embryonic day (E6)/ HH stage 29) (Fig. 1A). *TGIF1* qRT-PCR was performed on male and female gonads before (E4.5), during (E6.5) and after (E8.5) gonadal sex differentiation to validate sexually dimorphic expression. Quantitative RT-PCR showed a significant increase in *TGIF1* expression in female gonads from the onset of sexual differentiation (E6.5-E8.5) (Fig. 1B), consistent with the RNA-seq. *TGIF1* was also expressed in male gonads, but at consistently lower levels. *TGIF1* has a paralogue, *TGIF2*, with which it shares spatial and temporal expression in other developmental contexts (Shen and Walsh, 2005). *TGIF2* expression was not sexually dimorphic in the gonad RNA-seq data (Fig. S1A). This data was also confirmed by qRT-PCR (Fig. S1B).

**Fig. 1.**
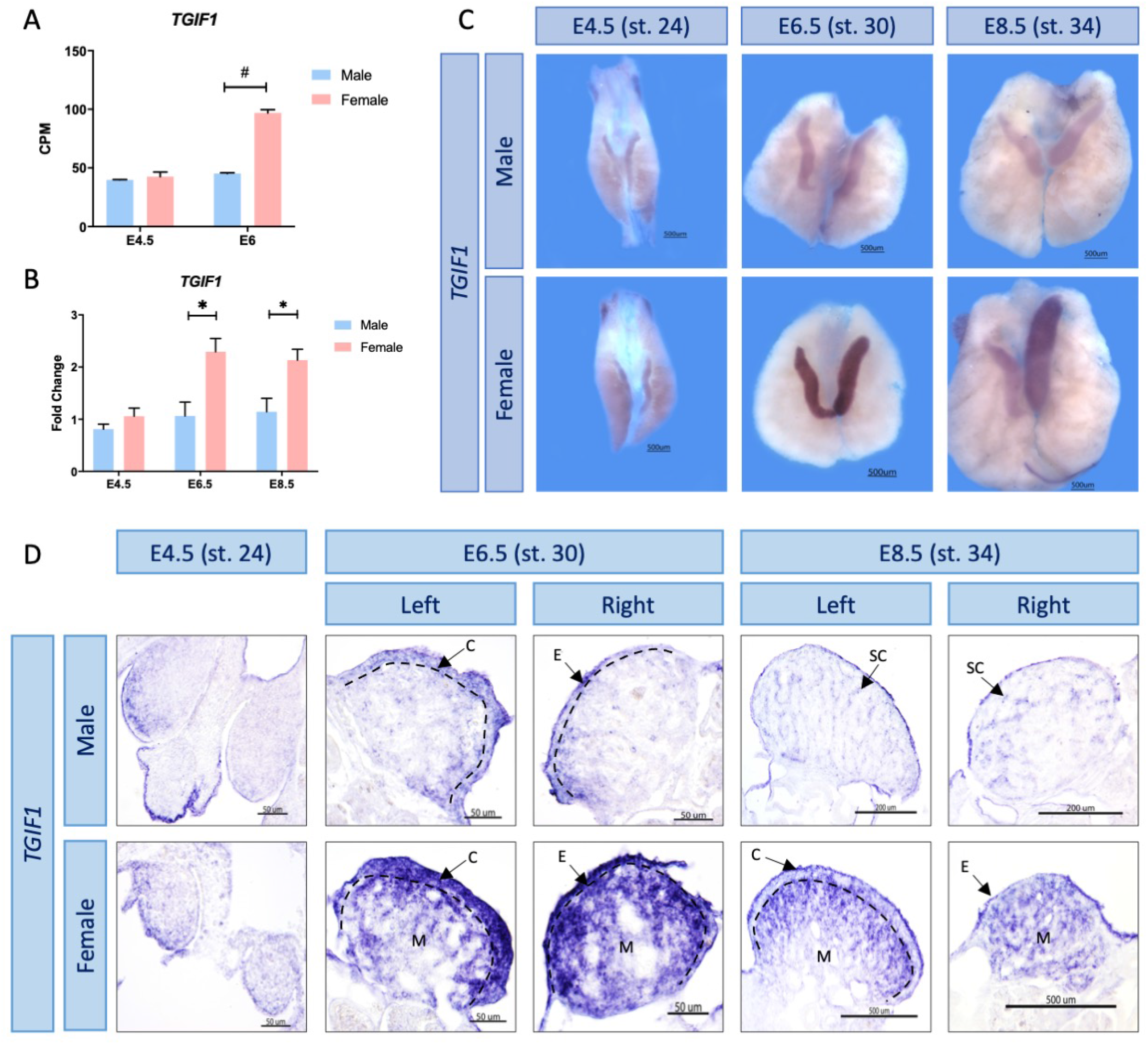
*TGIF1* expression profile in chicken gonads. (A) *TGIF1* gonadal RNA-seq mRNA expression levels in count per million (CPM) at blastoderm stage, before (E4.5) and on the onset (E6) of sex determination. # = false discovery rate (FDR) <0.001. (B) *TGIF1* gonadal mRNA was quantified by qRT-PCR. Expression level is relative to β-actin and normalized to E4.5 female. (Bars represent Mean ± SEM, n=6. * = adjusted p value <0.05. Multiple t-test and Holm-Sidak posttest. (C) *TGIF1* time course mRNA expression in embryonic chicken gonads, as assessed by whole mount in situ hybridization. (D) Sections of the *TGIF1* whole mounts in situ hybridizations. Arrows indicate the cortex (C), epithelium (E), medulla (M) and seminiferous cords (SC). Dashed black lines indicate the cortical-medulla limit.

### *TGIF1* is expressed in the ovarian cortical and medullary pre-granulosa cells

For spatial expression analysis of *TGIF1*, whole mount *in situ* hybridization was performed on male and female gonads at different developmental timepoints; before (E4.5/stage 24), during (E6.5/stage 30) and after (E8.5/ stage 34) sexual differentiation. *TGIF1* mRNA expression was stronger in female compared than male gonads from E6.5 / HH stage 30 (Fig. 1C), consistent with the RNA-seq and qRT-PCR results. In developing ovaries, *TGIF1* mRNA was localized in the cortex and in the medulla (Fig. 1D). In males, expression was detected at the surface epithelium at E6.5, though weaker than in the developing ovary (Fig. 1D). After the onset of sexual differentiation at E8.5 in males, weak expression was detected in the seminiferous cords of the medulla.

The developing chicken ovary comprises two distinct compartments: the outer cortex, which becomes thickened in the left ovary and is the site of oogenesis, and an inner medulla comprising interstitial, supporting and steroidogenic cells (Smith and Sinclair, 2004, Estermann et al., 2020). Co-localization with specific markers was performed to determine the cell types expressing *TGIF1* in the ovary. Aromatase and cytokeratin immunofluorescence were performed following *TGIF1* i*n situ* hybridization on tissue sections (Fig. 2). Aromatase marks estrogenic pre-granulosa cells, while cytokeratin marks cortical cells. In the left ovary, *TGIF1* was expressed in the cortical cells, colocalizing with cytokeratin. Lack of *TGIF1* expression in the right female gonad corresponded with the lack of a proliferating cortex. *TGIF1* mRNA also co-localized with the medullary pre-granulosa marker, aromatase, in both left and right gonads. *TGIF1* was not expressed in the interstitial cells between medullary cords nor in the juxtacortical medulla of female gonads (Fig. 2). In summary, *TGIF1* expression was restricted to cortical and pre-granulosa cells, both key cell types in ovarian development and differentiation.

**Fig. 2.**
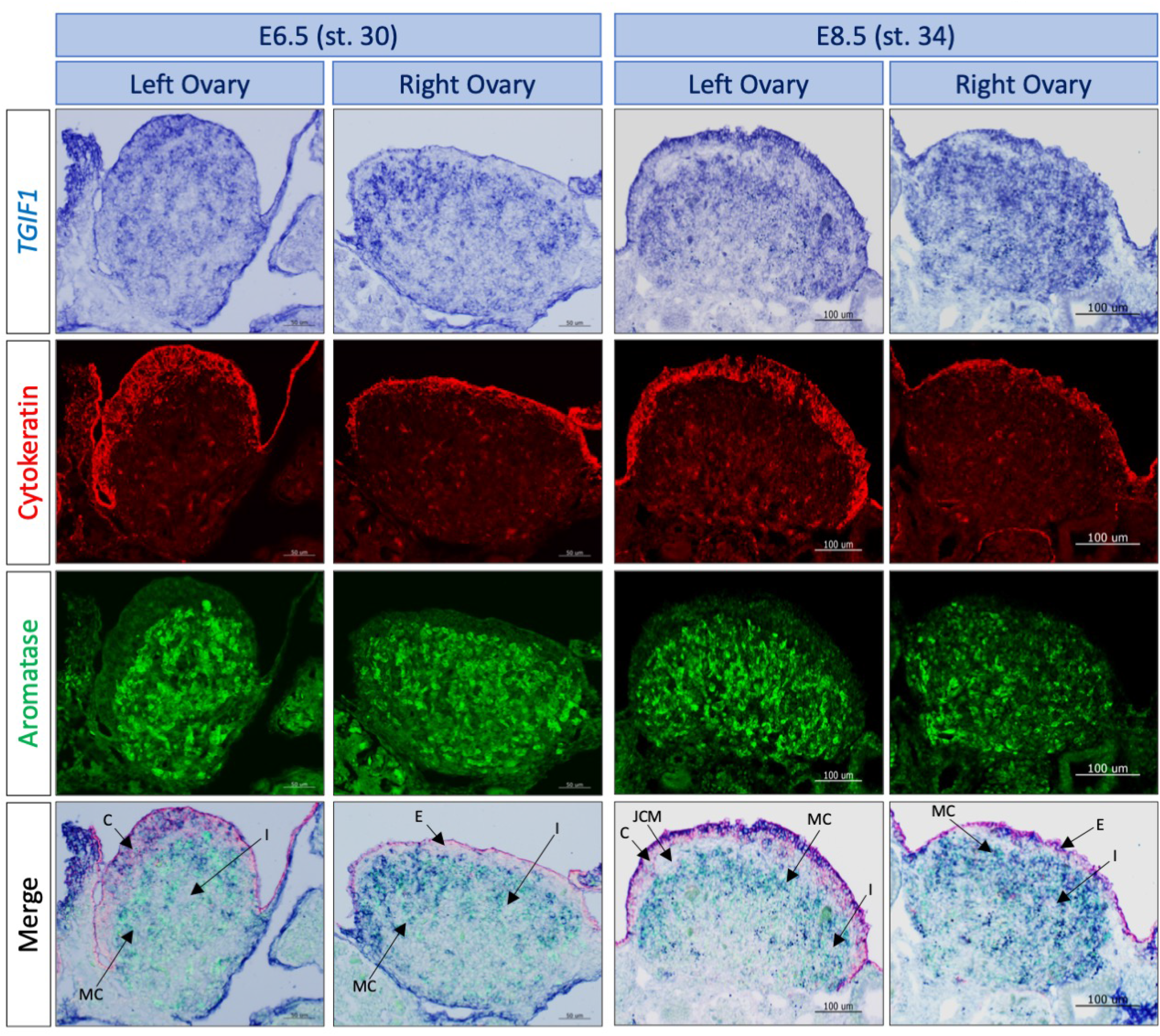
*TGIF1* expression colocalizes with key ovarian cells. *TGIF1* whole mount *in situ* hybridization was performed in E6.5 and E8.5 female urogenital systems. 10 um sections were processed for immunofluorescence against Cytokeratin (cortical cells marker) and Aromatase (pre-granulosa cells marker). *TGIF1* expression colocalize with both female markers in left ovaries. Arrows indicate the interstitial (I), cortical (C), medullary cords (MC), epithelial (E) and juxtacortical medulla (JCM) cells.

### *TGIF1* expression is sensitive to estrogens

Ovarian differentiation in birds is regulated by estrogen, catalyzed by the female restricted enzyme aromatase. *In ovo* injection of 17β-estradiol (E2) or the aromatase inhibitor fadrozole cause feminization and masculinization of the gonads, respectively (Bannister et al., 2011, Guioli et al., 2020). To determine whether *TGIF1* is responsive to estrogen signaling during ovarian development, sex reversal experiments were conducted. *TGIF1* was assayed following masculinization of female embryos with fadrozole, which inhibits aromatase enzyme action, or by applying estrogen to male embryos to induce feminization. *TGIF1 in situ* hybridization was performed in E9.5 male and female urogenital system (UGS) treated with 17-β-estradiol (E2) or vehicle (Control) at E3.5 (Fig. 3). Male gonads treated with E2 were morphologically feminized, with female-like asymmetry, characterized by a larger left and smaller right gonad. These gonads also showed structural organization typical of an ovary, with a thickened cortex (cytokeratin positive), aromatase positive pre-granulosa cells in the medulla and downregulated expression of the testis marker, anti-Müllerian hormone (AMH) (Fig. 3). *TGIF1* expression was upregulated in males treated with E2, compared with the vehicle control, showing a similar expression pattern to females (Fig. 3).

**Fig. 3.**
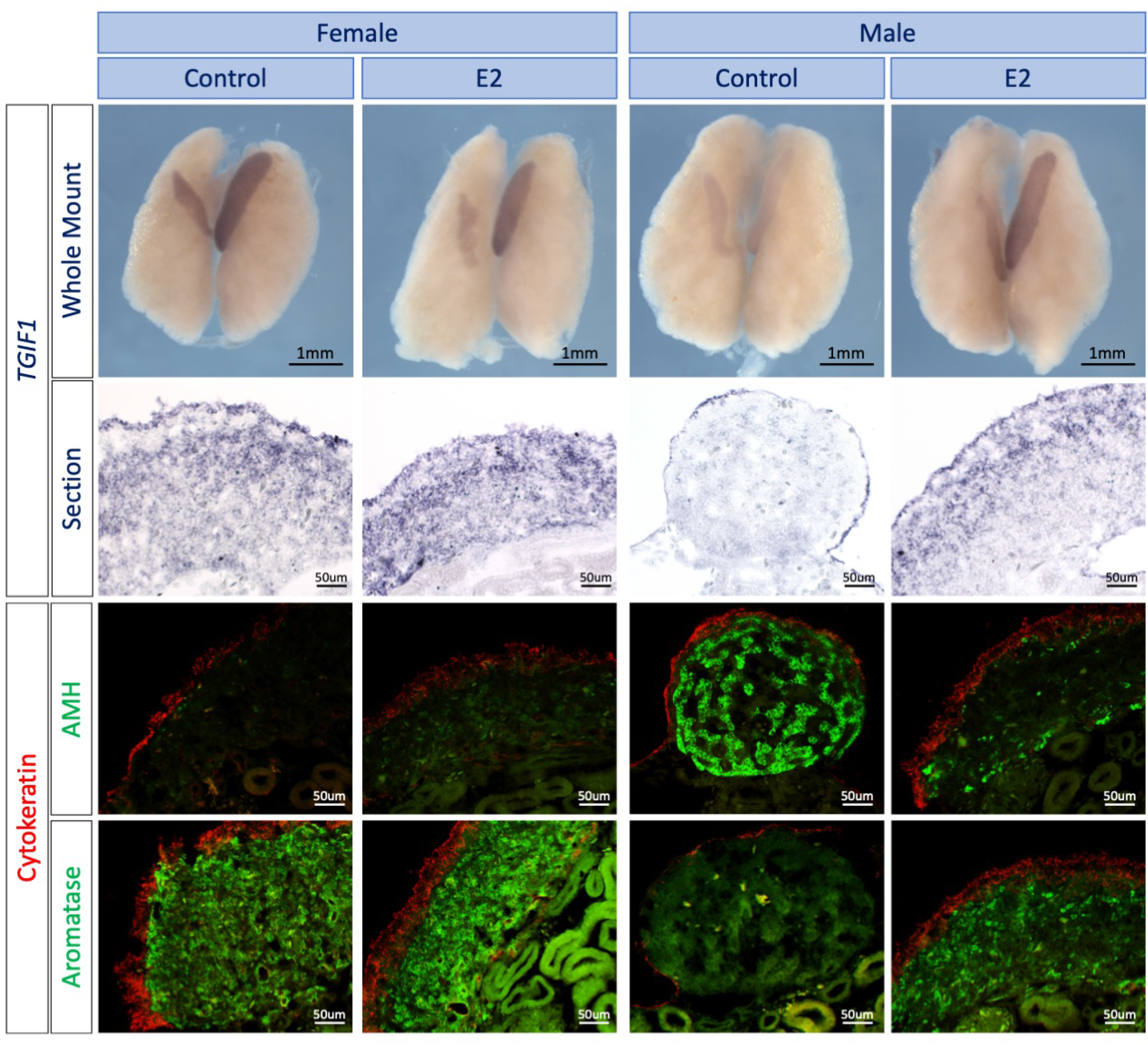
Estrogens induce *TGIF1* expression in ZZ gonads. *TGIF1* whole mount *in situ* hybridization was performed in E9.5 male and female urogenital systems treated *in ovo* with 17β-estradiol (E2) or vehicle (Control). Tissues were sectioned and immunofluorescence for aromatase (pre-granulosa marker) or AMH (Sertoli cell marker) and Cytokeratin (cortical marker) were performed to evaluate the efficacy of the sex reversal.

Female gonads treated with the aromatase inhibitor (AI) were masculinized, as expected. Female-type gonadal asymmetry was markedly reduced, and gonads showed testicular like morphology, containing AMH positive testicular cords, and a reduced cortex and reduced aromatase positive cells (Fig. 4A). *TGIF1* expression was also reduced in female gonads treated with aromatase inhibitor, consistent with the gonadal sex reversal (Fig. 4A). To quantify this change, *TGIF1* qRT-PCR was performed in E8.5 male and female gonads exposed to AI or vehicle (Control). Consistent with the *in situ* hybridization data, female gonads treated with AI showed a significant reduction of *TGIF1* expression in comparison with the vehicle control (Fig. 4B). Altogether, these results indicate that *TGIF1* mRNA expression responds to estrogens during ovarian differentiation in the chicken embryo.

**Fig. 4.**
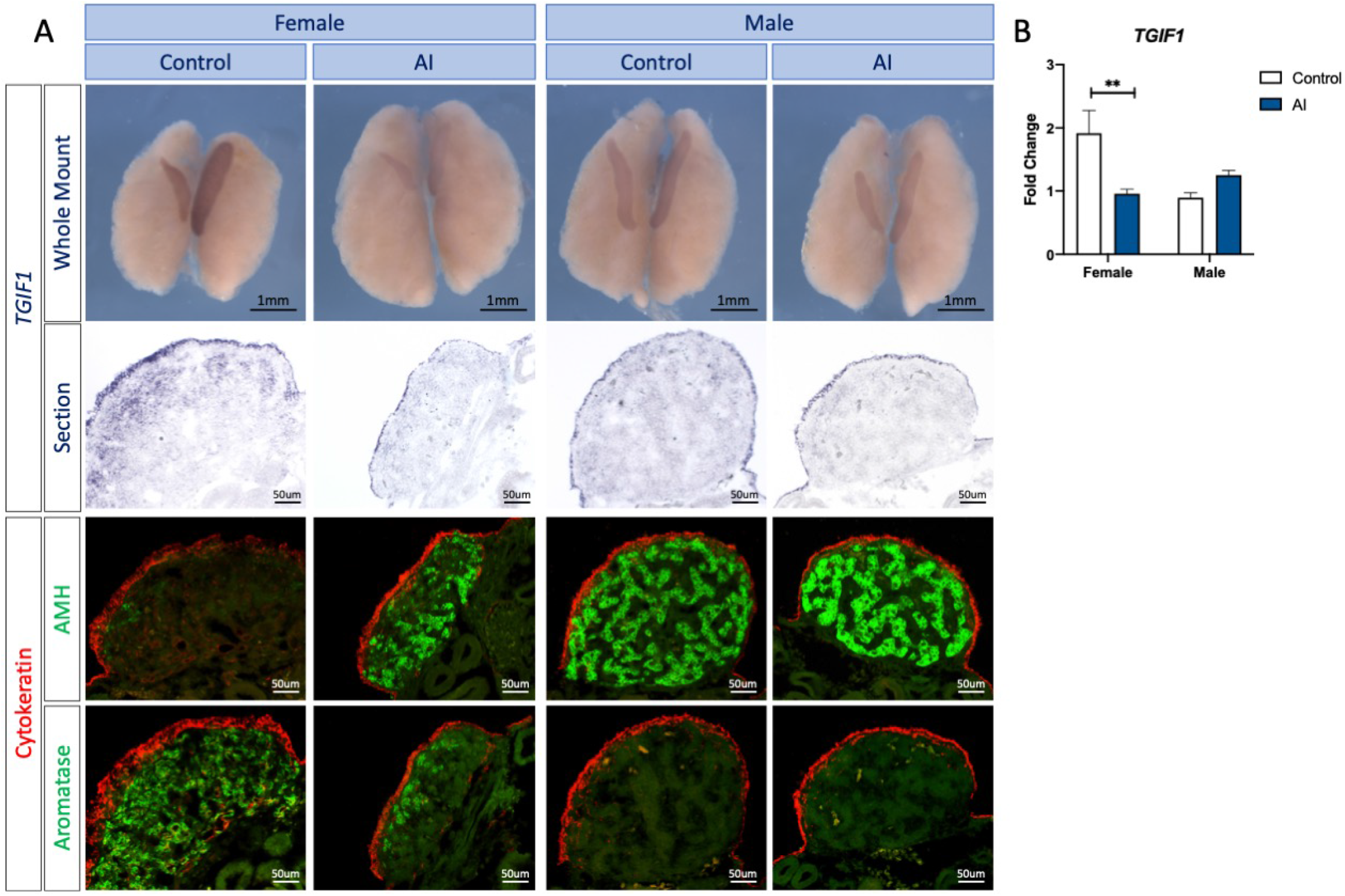
Estrogen synthesis inhibition by fadrozole results in downregulation of TGIF1 in ZW gonads. (A) *TGIF1* whole mount *in situ* hybridization was performed in E9.5 male and female urogenital systems treated *in ovo* with the aromatase inhibitor fadrozole (AI) or vehicle (Control). Tissues were sectioned and immunofluorescence for aromatase (pre-granulosa marker) or AMH (Sertoli cell marker) and Cytokeratin (cortical marker) were performed to evaluate the efficacy of the sex reversal. (B) *TGIF1* qRT-PCR was performed in gonadal samples of E9.5 embryos treated with the aromatase inhibitor (AI) or vehicle (Control). Expression level is relative to β-actin and normalized to male PBS. Bars represent Mean ± SEM, n=6. ** = adjusted p value <0.01. Multiple t-test and Holm-Sidak posttest.

### *TGIF1* over-expression in left testis results in gonadal feminization

To examine the effects of TGIF1 over-expression *in ovo*, electroporation of DNA constructs was used. Gonadal epithelial cells can be specifically targeted by performing electroporation of plasmid DNA into the coelomic epithelium at E2.5 without effecting the underlying medullary cord cell population (Estermann et al., 2020). This method allows insight into the role of TGIF1 specifically in the ovarian epithelium/cortex. TGIF1 open reading frame was cloned into TOL2-CAGGS-GPF vector, which, in presence of the transposase, integrates into the genome, stably expressing TGIF1 and GFP in the targeted and daughter cells (Sato et al., 2007). The ability of this construct to overexpress TGIF1 was assayed in vitro in the DF1 chicken fibroblastic cell line. Cells transfected with TGIF1 overexpressing plasmid significantly expressed around 30 times more TGIF1 than the empty plasmid (GFP control) (Fig. S1C).

TOL2-CAGGS-GFP-T2A-TGIF1 (TGIF1 OE) or TOL2-CAGGS-GFP (GFP Control) plasmid were co-electroporated with a plasmid expressing transposase into the left coelomic epithelium at E2.5 (stage 14). Embryos were collected at E8.5, sexed and immunofluorescence was performed against different gonadal markers. Figure 5 and 6 show the results of these experiments in which TGIF1 was over-expressed in the gonadal cortex of male embryos. In the absence of a suitable antibody to detect TGIF1 in chicken, GFP was used as a marker of electroporation. As expected, GFP was detected in the gonadal cortical/epithelial cells (Fig. 5), and in the interstitial cells that they generate, but not in supporting cells in males (Fig. 6) and females (Fig. S2). When *TGIF1* was over-expressed in female gonads, no structural or expression difference with the control was found (Fig. S2). In contrast, TGIF1 overexpression in left male gonads resulted in a change in the epithelial cell structure (cytokeratin positive, fibronectin negative), resembling cuboidal rather than squamous epithelium that is typical of the testis (Fig. 5A). Image quantification analysis indicated that TGIF1 over-expression resulted in a significant increase of the gonadal epithelial area (Fig. 5C) and the thickness of the epithelium (cell height) (Fig. 5C). In addition, an increment of cytokeratin positive mesenchymal cells was detected, suggesting augmentation of an epithelial to mesenchyme transition (EMT) (Fig. 5A). In male gonads overexpressing *TGIF1*, interstitial cells derived from the coelomic epithelial cells by EMT (GFP^+^, fibronectin^+^) accumulated underneath the epithelial layer, forming dense clusters (Fig. 6A). This accumulation of interstitial cells resembles the organization of the ovarian juxtacortical medulla (JCM) and resulted in a displacement of the testicular cords (AMH^+^) towards the more basal region of the gonad (Fig. 6B). Image quantification analysis indicated that TGIF1 over-expression resulted in a significant increase in the juxtacortical medulla area, compared with the controls (Fig. 6C).

**Fig. 5.**
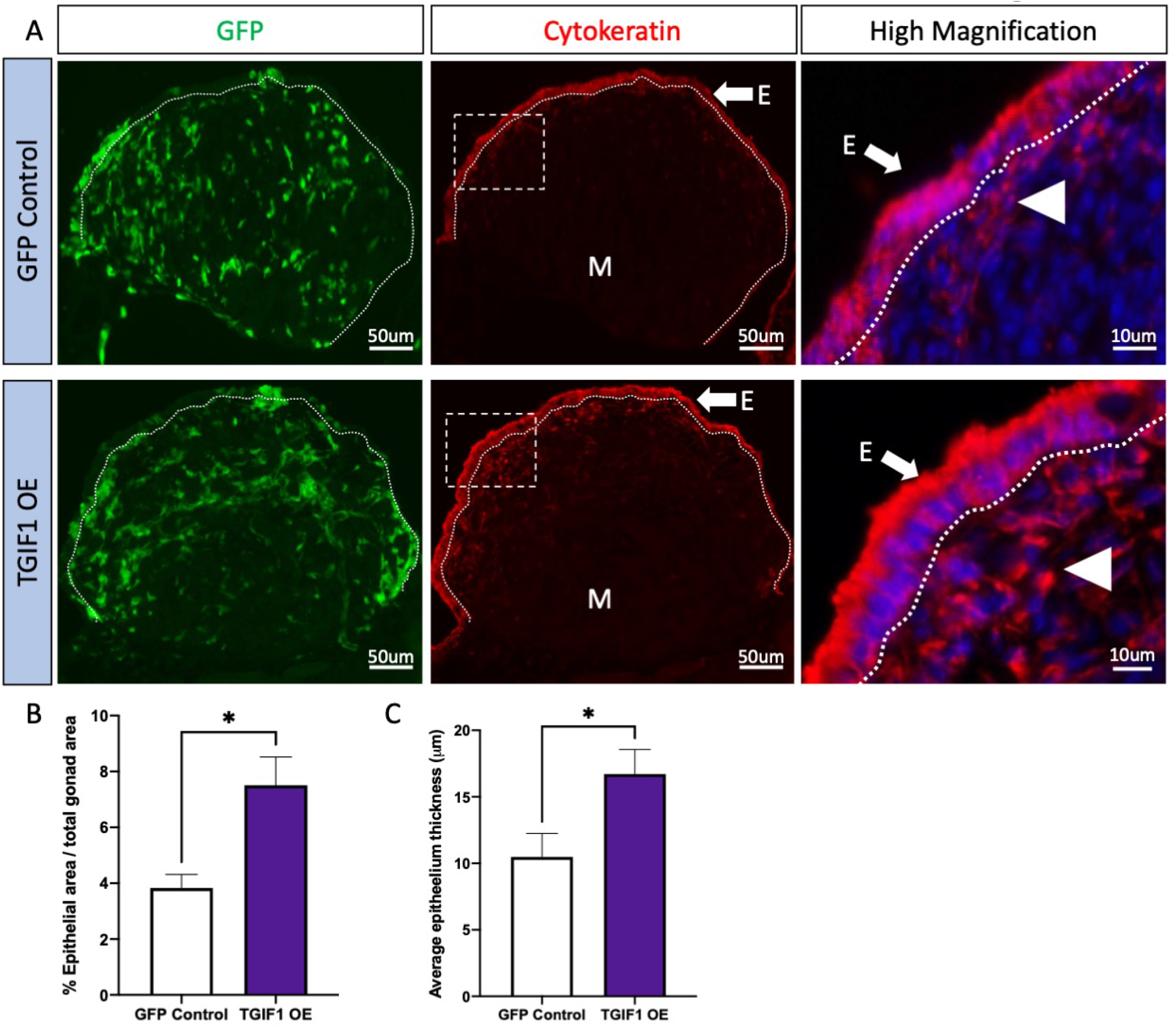
TGIF1 overexpression results in epithelial thickening. TOL2 TGIF1 overexpression (TGIF1 OE) or control (GFP Control) plasmids were electroporated in male left E2.5 coelomic epithelium. Gonads were examined at E8.5. (A) Immunofluorescence against cytokeratin (epithelial/cortical marker) were performed in transverse sections. Dashed box indicates the magnified area. Dotted line delineates the gonadal epithelium. White arrow indicates the epithelium (E), white arrowhead indicates EMT derived interstitial cell and M indicates medulla. (B) Quantification of the percentage of epithelial area, related to the total gonadal area in male gonads. (C) Quantification of the average epithelium thickness (in um) in control or TGIF1 overexpressing male gonads. Bars represent Mean ± SEM, n≥6. Unpaired two-tailed t-test. * = p value <0.05.

**Fig. 6.**
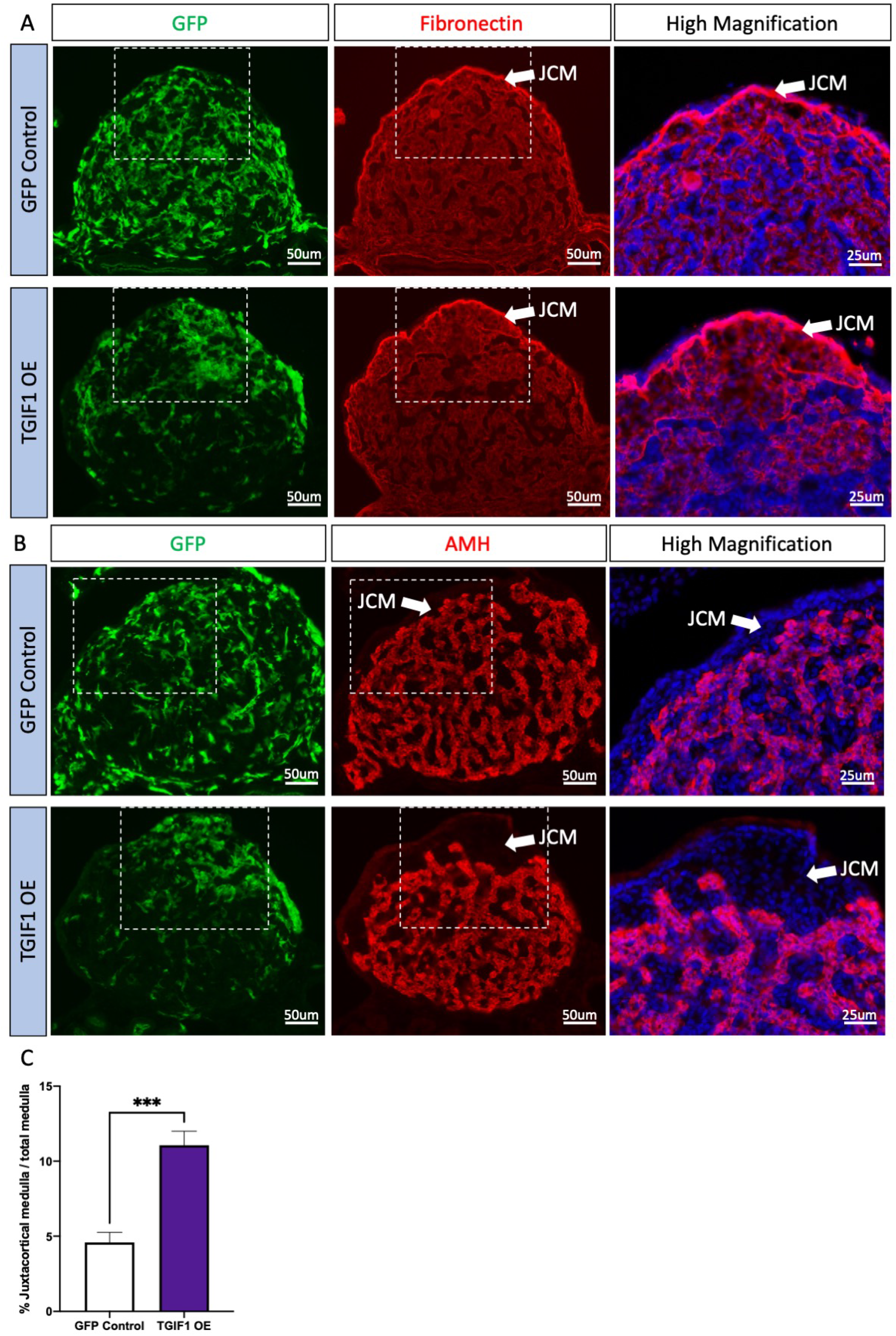
TGIF1 overexpression results in gonadal feminization. TOL2 TGIF1 overexpression (TGIF1 OE) or control (GFP Control) plasmids were electroporated in male left E2.5 coelomic epithelium. Gonads were examined at E8.5. Immunofluorescence against (A) fibronectin (interstitial cell marker) and (B) AMH (Sertoli cell marker) were performed in transverse sections. Dashed box indicates the magnified area. Dotted line delineates the gonadal epithelium. Arrows indicate the juxtacortical medulla (JCM). (C) Quantification of the percentage of juxtacortical medulla area, related to the total medullar area. Bars represent Mean ± SEM, n≥7. Unpaired two-tailed t-test. *** = p value <0.001.

To examine its effects on the gonadal medulla, TGIF1-GFP was over-expressed using the RCASBP viral vector. Unlike TOL2, this vector can spread horizontally to neighboring cells. This is important, as it can deliver transgenic expression to the medullary cord population which cannot be targeted by TOL2 electroporation. *TGIF1* ORF was cloned into RCAS(A)-GFP viral vector. RCAS(A)-GFP-T2A-TGIF1 (TGIF1 OE) or RCAS(A)-GFP (GFP Control) plasmids were electroporated in E2.5 left coelomic epithelium. Urogenital systems were collected at E7.5-E8.5, sexed and immunofluorescence was performed against different gonadal markers. Over-expression of TGIF1 in male gonads using this approach did not alter the testicular development. Supporting cells developed normally, AMH, SOX9 and DMRT1 expression was similar than the control gonads (Fig. S3), and no female markers (aromatase or FOXL2) were detected (data not shown). Instead, the same morphological changes were detected when TGIF1 was overexpressed only in the coelomic epithelial cells: displacement of the supporting cells from the sub-epithelial region (Fig. S3A and B) and an increased thickness of the gonadal surface epithelium, marked by diagnostic cytokeratin staining. The cells of the epithelium adopted a female-like cuboidal morphology instead of the squamous epithelium typical of the testis (Fig. 6). Altogether, this data suggests that TGIF1 miss-expression does not impact Sertoli cell differentiation in chicken gonads.

### Ovarian TGIF1 knock down inhibits gonadal cortex and juxtacortical medulla formation

*In ovo* gene knockdown was performed to assess the role of TGIF1 during cortical and juxtacortical medulla formation in the developing ovary. Four different shRNAs were designed against the *TGIF1* open reading frame and were cloned into the retroviral vector RCASBP (D) carrying a blue fluorescent protein (BFP) reporter. These were screened for knockdown efficiency *in vitro* using the DF1 chicken fibroblastic cell line. DF1 cells were transfected with plasmids carrying BFP-T2A and a non-silencing control shRNA (NS shRNA), or one of four different shRNAs designed for TGIF1 knockdown. After all cells became BFP positive, they were transfected with TOL2 plasmid over-expressing chicken TGIF1-GFP (TOL2-GFP-T2A-TGIF1). Plasmid expressing mCherry was used as a transfection control. 48 hours post transfection, cells were fixed, and GFP fluorescence was quantified as a measure of *TGIF1* knockdown (Fig. 8A). All of the shRNAs showed a significant decrease of GFP intensity. Sh988 showed the strongest inhibition (66%), followed by sh364 (57%), sh416 (46%) and sh318 (16%). These values were calculated using the mean of each group using NS shRNA as a control (100%) (Fig. 8B).

**Fig. 7.**
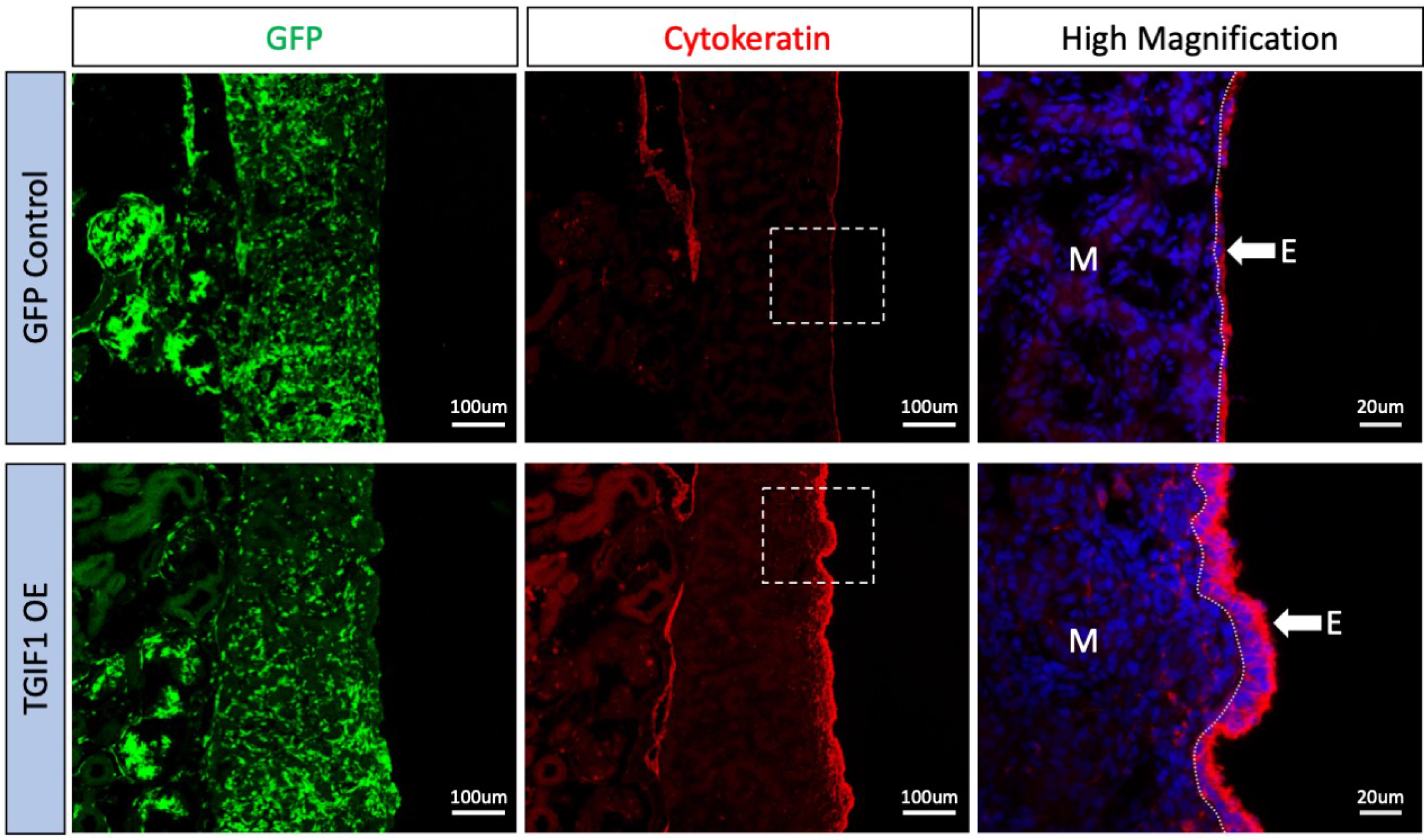
TGIF1 RCAS overexpression mimics the results from TGIF1 TOL2 overexpression. RCAS(A) TGIF1 overexpression (TGIF1 OE) or control (GFP OE) plasmids were electroporated in male left E2.5 coelomic epithelium. Male gonads were examined at E8.5 and immunofluorescence against cytokeratin (epithelial/cortical marker) was performed in longitudinal sections. Dashed box indicates the magnified area. Dotted line delineates the gonadal epithelium. White arrows indicate the gonadal epithelium (E). M indicates the medulla.

**Fig. 8.**
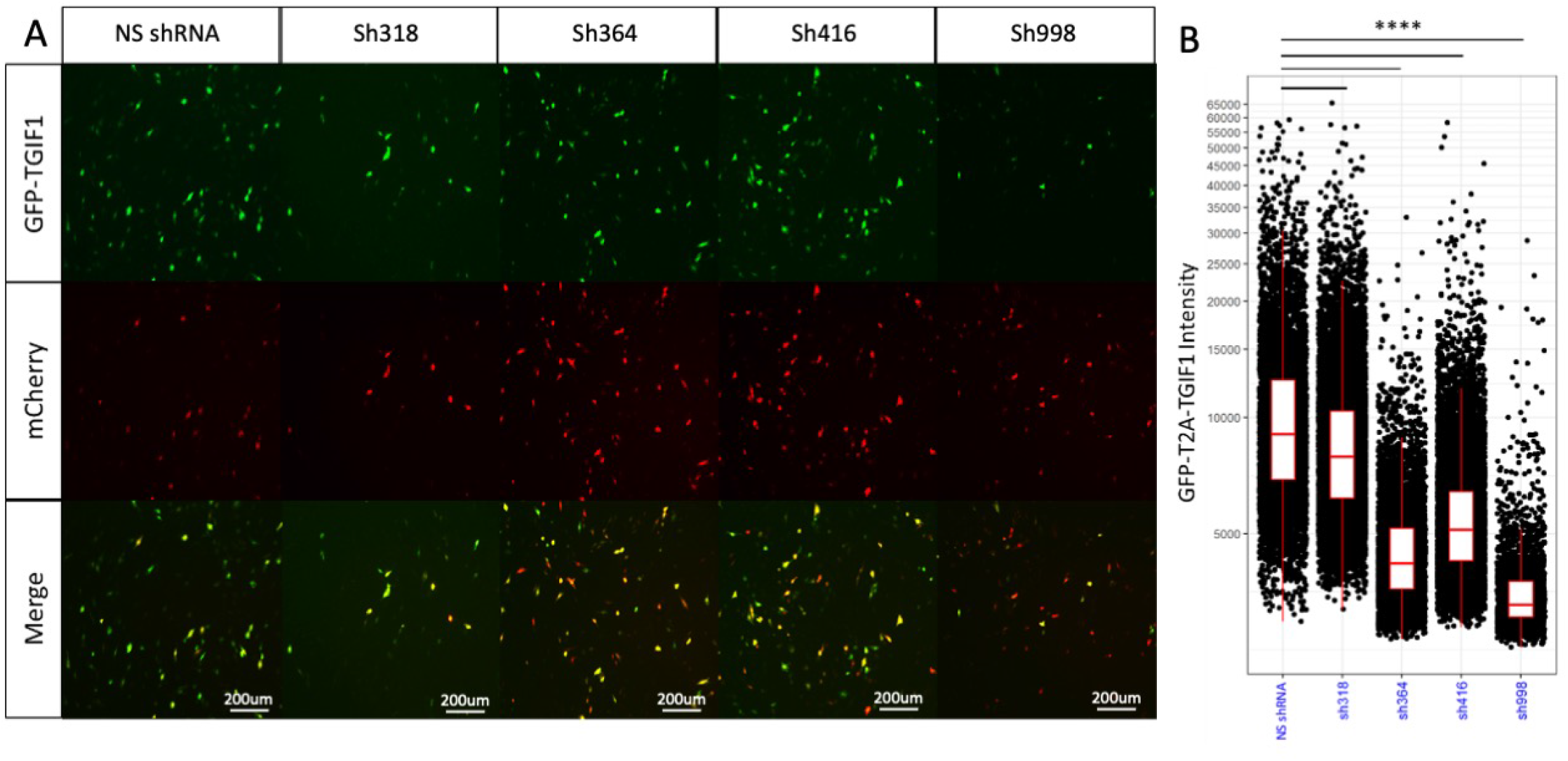
TGIF1 sh998 showed the higher repression of TGIF1 in vitro. Chicken DF-1 cells were transfected with a self-replicative viral plasmid containing BFP-T2A and a non-specific shRNA (NS shRNA) or with 4 different putative shRNA designed for TGIF1 knockdown (sh318, sh364, sh416 or sh998). After all cells were BFP positive, they were transfected with the overexpression TOL2-GFP-T2A-TGIF1 plasmid and a plasmid expressing mCherry (as a transfection control). After 48 hours cells were fixed and analyzed under the microscope. (A) Representative fluorescence images of the outcomes. (B) Imaris analysis of the GFP-T2A-TGIF intensity in mCherry positive (transfected) cells. Box plots show each sample’s median, interquartile ranges (IQR) and the whiskers extend to the highest/lowest value within 1.5 x IQR. Each dot represents an individual cell. t-test was performed using NS shRNA as a control condition. Dunnett’s multiple comparisons test was used as posttest. ****, p < 0.0001.

TGIF1 sh998 was cloned into a TOL2 vector expressing nuclear BFP. TOL2-TGIF1sh998-nBFP or TOL2-NSshRNA-nBFP were *in ovo* co-electroporated with TOL2-ACAGS-GFP (electroporation reporter) and transposase expressing plasmid into the left coelomic epithelium at E2.5. Urogenital systems were collected at E8.5, genetically sexed by PCR and immunofluorescence was performed for different gonadal markers. *TGIF1* knock down resulted in a substantial size reduction of the targeted left ovaries, in comparison with the controls electroporated with NS shRNA. Aromatase positive pre-granulosa cells were still present in the gonadal medulla (Fig. 9A) and no Sertoli cell markers (SOX9, AMH, DMRT1) were up-regulated (data not shown). Strikingly, an ovarian cortex was absent in the female *TGIF1* knock down gonads, as reveled by cytokeratin expression (Fig. 9B). Instead, the epithelial cells exhibited a flattened morphology similar to the right gonadal epithelium or the testicular epithelium (Fig. 9B). In addition, these ovaries lacked a clear juxtacortical medulla (JCM), evidenced by the absence of condensed fibronectin positive cells in between the epithelium and the aromatase positive pre-granulosa cells (Fig. 9C). Due to the absence of a defined cortex, the germ cells remained in the medulla (similar to their fate in the right gonad) (Fig. 9D). Altogether, these results indicate that TGIF1 is necessary to develop an ovarian cortex. Moreover, TGIF1 is the first gene reported to be required for the juxtacortical medulla formation.

**Fig. 9.**
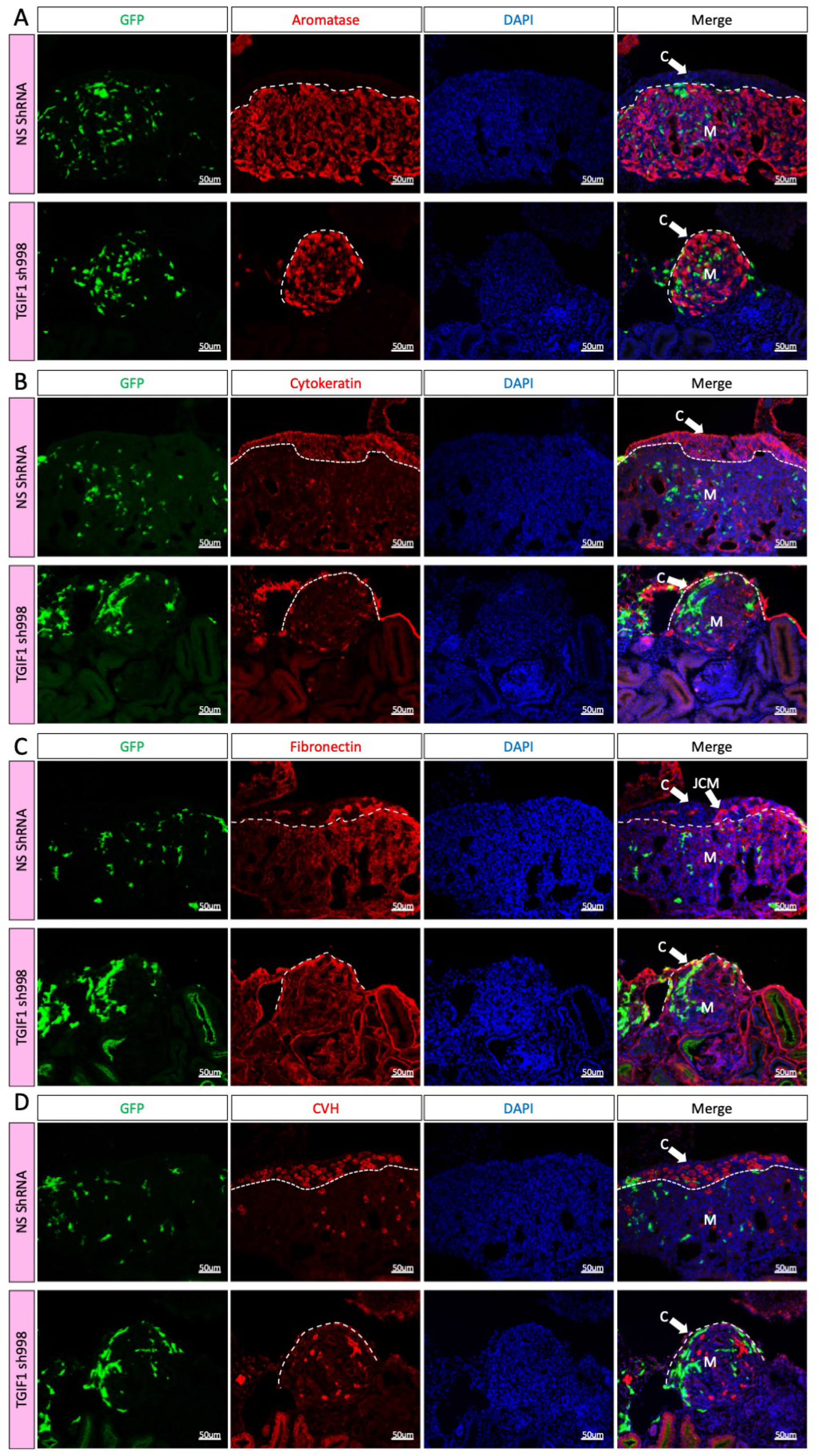
TGIF1 Knock down ablates cortical and juxtacortical medulla formation in female gonads. TOL2 TGIF1 knock down (TGIF1 sh998) or non-silencing control (NS shRNA) plasmids were co-electroporated with a GFP expressing plasmid (reporter) in female left E2.5 coelomic epithelium. Gonads were examined at E8.5. Immunofluorescence against (A) aromatase (pre-granulosa marker), (B) cytokeratin (epithelial/cortical marker), (C) fibronectin (interstitial cell marker) and (D) CVH (germ cell marker) were performed in transverse sections. Dashed line delineates the gonadal epithelium. Arrows indicate the cortex (C) or the juxtacortical medulla (JCM). M indicates the medulla.

## DISCUSSION

Gonadal sex differentiation provides an ideal model for studying progressive cell fate decisions (Lin and Capel, 2015, Munger and Capel, 2012). During gonadal morphogenesis, cell lineages differentiate into ovarian or testicular cell types. The first cells to differentiate are the supporting cells (Sertoli cells in males, pre-granulosa cells in females) (Nef et al., 2019). These cells then signal to control differentiation of the other gonadal lineages, including steroidogenic and non-steroidogenic cells, and they also influence germ cell fate (spermatogonia in males, oogonia in females) (Stevant et al., 2019, Stevant and Nef, 2019). Due to the central role of the supporting cell population, most research in the field has focused on the granulosa vs Sertoli cell fate decision. Less is understood about the development and role of the gonadal surface (coelomic) epithelium. However, lineage tracing and single-cell RNA-seq have shown that this gonadal compartment has significantly different roles in mammalian versus avian models. In the mouse embryo, the surface epithelium is central to gonadal differentiation. This layer of cells is the source of most somatic cell progenitors in the embryonic murine gonad (DeFalco et al., 2011, Nicol and Yao, 2014). The surface epithelial cells express the transcription factors Wt1, Gata4 and Sf1, they proliferate to first give rise first to the supporting cell lineage, and, subsequently, at least some of the steroidogenic population (Hatano et al., 2010, Karl and Capel, 1998, Stevant and Nef, 2019, Stevant et al., 2019, Nef et al., 2019). Somatic cells of the surface epithelium in mouse divide asymmetrically, producing one daughter cell that remains at the surface and one that undergoes an epithelial to mesenchyme transition (EMT), ingressing into the gonad. This process is regulated by Notch signaling, via the antagonist, Numb (Lin et al., 2017). Homeobox transcription factors such as Emx2, Six1 and Six4 contribute to this EMT (Kusaka et al., 2010, Fujimoto et al., 2013). In contrast to mouse, lineage tracing in the chicken embryo clearly shows that proliferating surface epithelium gives rise to non-steroidogenic interstitial cells, not the supporting cell lineage (which derives from mesonephric mesenchyme) (Sekido and Lovell-Badge, 2007, Estermann et al., 2020). Furthermore, epithelial cells differentiate into a stratified layer of cortical cells in the left female gonad, whereas this process does not occur in males (Estermann et al., 2020). The development of a thickened left gonadal cortex is critical for proper ovary formation and female reproduction in birds. Germ cells accumulate in the ovarian cortex during embryonic stages and are signaled to enter meiotic prophase (Smith et al., 2008). After hatching, germ cells development proceeds as the cortex is the site of folliculogenesis in avians (Johnson and Woods, 2009, Li et al., 2016, Hu et al., 2021). The importance of the cortex is revealed by the asymmetry of female avian gonadal development. The right gonad fails to elaborate as cortex in females and germ cells remain in the medullar, where they eventually become atretic (Guioli et al., 2014).

The results presented here demonstrate that the TALE homeobox gene, *TGIF1*, plays a key role in development of the gonadal cortex in the chicken embryo. This gene is upregulated during female but not male gonads development (Fig. 1). It is strongly expressed in the female gonadal surface epithelium at onset of sexual differentiation (E6.5/stage 30), and in the gonadal medulla. Targeted over-expression in the male surface epithelium induces a thickened cortex, while targeted knockdown in in females blocks proper cortical layer development. Manipulation of expression in the medulla did not have an overt effect upon gonadogenesis. Furthermore, *TGIF1* expression was responsive to modulation of estrogen, which is essential for ovarian development in birds (Scheib, 1983, Elbrecht and Smith, 1992, Vaillant et al., 2001a). Inhibition of the estrogen-synthesizing enzyme, aromatase, resulted in down-regulation of *TGIF1* expression (Fig. 4). This indicates that TGIF1 is a downstream target of estrogen, either directly or indirectly, during ovary formation. In the chicken embryo, two roles are ascribed to the estrogen that is synthesized by medullary cords cells of female embryos at the onset of gonadal sex differentiation. Firstly, estrogen acts in the medulla itself to antagonize the induction of the testis factors, DMRT1 and SOX9 (Smith et al., 2003, Ioannidis et al., 2021). Secondly, acts on the surface epithelium in a paracrine fashion, where it stimulates development of the gonadal cortex (Gasc and Stumpf, 1981, Wartenberg et al., 1992). Correspondly, estrogen receptor α (ER-α), is expressed in both gonadal compartments in chicken embryos (Andrews et al., 1997, Gonzalez-Moran, 2014.) Exogenous estrogens can induce cortical cell differentiation in embryonic male (ZZ) gonads (Guioli et al., 2020). While gonadal asymmetry in the chicken is driven by asymmetric expression of *Pitx2* in the cortex (Rodriguez-Leon et al., 2008, Guioli and Lovell-Badge, 2007), cortical cell proliferation is related to estrogen action. ER-α is expressed in the left but not the right gonadal epithelium. This is consistent with the cortical development in the left but not in the right gonad. RNAi or domain negative-mediated downregulation of ER-α cause a reduction in cortical size, indicating that ER-α and estrogens are essential for cortex formation (Guioli et al., 2020). Here, we found that *TGIF1* expression was induced by estrogen and downregulated when estrogen synthesis was inhibited (Fig. 3 and 4). Moreover, *TGIF1* and ER-α are both expressed the left gonadal epithelium (and in supporting cells of the medulla), suggesting that estrogens, through ERα, could regulate *TGIF1* expression in chicken ovaries. This would also explain why *TGIF1* is expressed in the left gonadal epithelium but not in the right (Fig. 2). Similar to ERα, TGIF1 knock down in female gonads resulted in lack of cortical development, despite the presence of estrogens (aromatase expression was not perturbed). This indicates that *TGIF1* is required for ovarian cortical formation, acting downstream of the estrogen signaling pathway. It will be of interest to examine the regulatory region of the chicken TGIF1 gene for estrogen response elements.

The data presented here indicate that one of the functions of TGIF1 is the maintenance of columnar epithelial cells the gonadal cortex. The epithelium in the left and right gonads in both males and female chicken embryos at E6.5 shows an asymmetry, being thicker in the left than the right gonad (Guioli et al., 2014). In females this structure continues proliferating, whereas in the male, it flattens and to a squamous monolayer. *TGIF1* over-expression in males did not induce the formation of a multilayered female like cortex. However, epithelial cell thickness was increased (Fig. 5 and 6). This suggests that TGIF1 is required for maintaining columnar epithelial structure and inhibiting a squamous phenotype. *Tgif1*/*Tgif2* double null mouse embryos display disorganized epiblasts and lacked the typical columnar epithelial morphology (Powers et al., 2010). This suggests that the role of TGIF1 in maintaining the epithelium structure may be a conserved function during embryogenesis.

TGIF1 May be acting through a number of mechanisms to promote development of the ovarian cortex in the chicken embryo. *TGIF1* encodes a homeodomain transcription factor of the TALE family (Three Amino Loop Extension). At least three signaling pathways have been linked to TGIF1 function; TGF (Wotton et al., 1999b), retinoic acid (Bertolino et al., 1995) and Wnt/β-catenin (Zhang et al., 2015b). All of these pathways are known to be engaged in the embryonic gonads in chicken and in mouse. TGIF1 is a TGF-β signaling inhibitor, binding to phospho-Smad2, recruiting histone deacetylases and acting as a co-repressor of Smad target genes (Wotton et al., 1999a). Chicken ovaries exposed to TGF-β1 display a reduction of somatic cells due to decreased cell proliferation (Mendez et al., 2006). In addition, there is a reduction in the number of germ cells in the cortex and an increased number in the medulla (Mendez et al., 2006).This suggests an effect in the cortical compartment or in the capacity of germ cells to migrate. In mice, nodal, activin and TGF-β signalize through Smad 2/3/4 and are key in testicular development, suppressing the pre-granulosa program (Gustin et al., 2016, Wu et al., 2013). In contrast, BMP molecules such as BMP2 signal through Smad 1/5/8 and are important for ovarian differentiation is mouse (Kashimada et al., 2011). TGIF1, being expressed in the female supporting cells and cortex, could act to repress the Smad 2/3 masculinizing signaling and allowing BMPs to induce ovarian differentiation. Further research should explore the role of TGIF1 modulating the balance between BMP vs TGF-β pathways in the gonadal context.

In chicken, lineage tracing experiments show that non-steroidogenic interstitial cells derive from the gonadal surface epithelium by an EMT (Estermann et al., 2020). The ultimate fate of these cells is poorly known, but they likely contribute to peritubular myoid cells in males but their role in females is obscure. However, in the chicken, there is an accumulation of interstitial cells directly underneath the cortex in females, forming a zone called the so-called juxtacortical medulla. This structure is not present in testis, and its functional significance is not known. However, several genes show restricted expression in the JCM later in development, such as CYP26B1, responsible for retinoic acid degradation (Smith et al., 2008). Recently, TGIF1 was found to be expressed in chicken dorsal neural tube and in delaminating cardiac neural crest, where it is required for the formation of mesenchymal derivatives of the crest (Gandhi et al., 2020). In addition, TGIF1 is associated with increased breast, lung and colorectal cancer migration and metastasis (Haider et al., 2020, Xiang et al., 2015, Wang et al., 2017) and can activate the Wnt/β-catenin signaling to promote cancer cell proliferation and migration (Wang et al., 2017, Zhang et al., 2015b). TGIF1 over-expression in male coelomic epithelium induced fibronectin positive interstitial cells accumulation underneath the gonadal epithelium, despite lacking a fully developed cortex (Fig. 5B). This process would appear to be independent estrogen signaling, due to the absence of aromatase expression, and consequently, estrogens in the male gonads miss-expressing TGIF1. Consistently, ovaries lacked a juxtacortical medulla when TGIF1 was knocked down (Fig. 9). This indicates that TGIF1 induces EMT in the epithelial cells to generate interstitial cells, which accumulate beneath the gonadal epithelium. TGIF1 is the first gene reported to be required for the juxtacortical medulla formation. Interestingly, TGIF1 is expressed in the epithelial cells but not in the interstitial cells, suggesting that TGIF1 is downregulated after the EMT, consistent with its role of maintaining the surface epithelium.

TGIF1 was also found to be expressed in the pre-granulosa cells, colocalizing with aromatase (Fig. 2). This suggest that TGIF1 could play a role in supporting cell differentiation. SOX9 is a marker of Sertoli cell, and it is known to have a role in repressing the female differentiation pathway and inducing and maintaining the male genetic program. When TGIF1 was over-expressed in mouse limb mesodermal micromass cultures, chondrogenic markers, such as Sox9, were downregulated (Lorda-Diez et al., 2009). When TGIF1 was silenced, SOX9 expression was up-regulated, suggesting a direct or indirect role of TGIF1 in inhibiting SOX9 (Lorda-Diez et al., 2009). Here, over-expressing TGIF1 in testicular supporting cells did not result in a reduction of SOX9 expression or the upregulation of pre-granulosa markers (Fig. S3). This suggest that this role of TGIF1 is not conserved among mouse and chicken or that its function differs between limbs and gonads. The current data presented here indicate that TGIF1, by itself, has no role in early differentiation of supporting cells in the chicken model.

TGIF1 and TGIF2 share similar spatial and temporal expression during embryonic development. In addition, they have similar binding domains, suggesting functional redundancy (Shen and Walsh, 2005, Powers et al., 2010). TGIF2 has redundant functions with TGIF1, but they need to be co-expressed in the same cells in order to have a compensatory effect (Lee et al., 2015). In chicken gonads, both TGIF1 and TGIF2 are expressed in the gonads, but RNA-seq data showed that TGIF2 expression is lower than TGIF1 (Fig. 1A & S1A). In ovarian TGIF1 knock down experiments, TGIF2 expression levels were not able to rescue the cortical and juxtacortical formation. This suggest that in chicken gonads TGIF1 and TGIF2 do not share the same functions, or they are not expressed in the same cell types. While TGIF1 not previously been linked to vertebrate gonadal development, *Drosophila* TGIF and tammar wallaby TGIF2 are important for spermatogenesis (Ayyar et al., 2003, Hu et al., 2011). In chicken, TGIF2 expression was not sexually dimorphic. Due to the reported role in spermatogenesis, it would be interesting to study TGIF2 expression in adult birds to identify if this function is conserved among species.

In summary, in chicken ovaries, the data presented here indicated that activation of the ERα signaling pathway by estrogens induces the expression of TGIF1 in the gonadal epithelium of the female the chicken embryo. TGIF1 expression supports development of the ovarian cortex, inhibiting the epithelial flattening, and it induces formation of the juxtacortical medulla by increased EMT (Fig. 10). In chicken testis, TGIF1 expression is not induced in the gonadal epithelial cells due to the lack of estrogens and, consequently, ERα signaling. This results in the epithelial flattening, inhibiting the formation of the juxtacortical medulla (Fig. 10). Our results support the proposal that supporting cell differentiation and cortical sex differentiation are two independent processes (Guioli et al., 2020, Ioannidis et al., 2021). This research introduces TGIF1 as one of the main regulators of cortical differentiation. In addition, we identified TGIF1 as the first known regulator required for juxtacortical medulla formation and provide evidence that a fully developed cortex or estrogens are not required for this process. Future research should focus on the downstream targets of TGIF1 in regulating this process. In addition of its role as a transcription factor, TGIF1 was also associated with several functions in signaling pathways. These includes TGF-β, retinoic acid and WNT/β-catenin pathways (Lorda-Diez et al., 2009, Liu et al., 2014, Gongal and Waskiewicz, 2008, Castillo et al., 2010, Zhang et al., 2015b). A comprehensive analysis of the role of TGIF1 in regulating cell signaling in the gonadal context is required to fully understand its role in gonadogenesis. This could also shed light in the role of TGIF1 in the supporting cell, which still remains unknown. Our research provides new insights in chicken ovarian differentiation and development, specifically in the process of cortical and juxtacortical medulla formation, a less explored field.

**Fig. 10.**
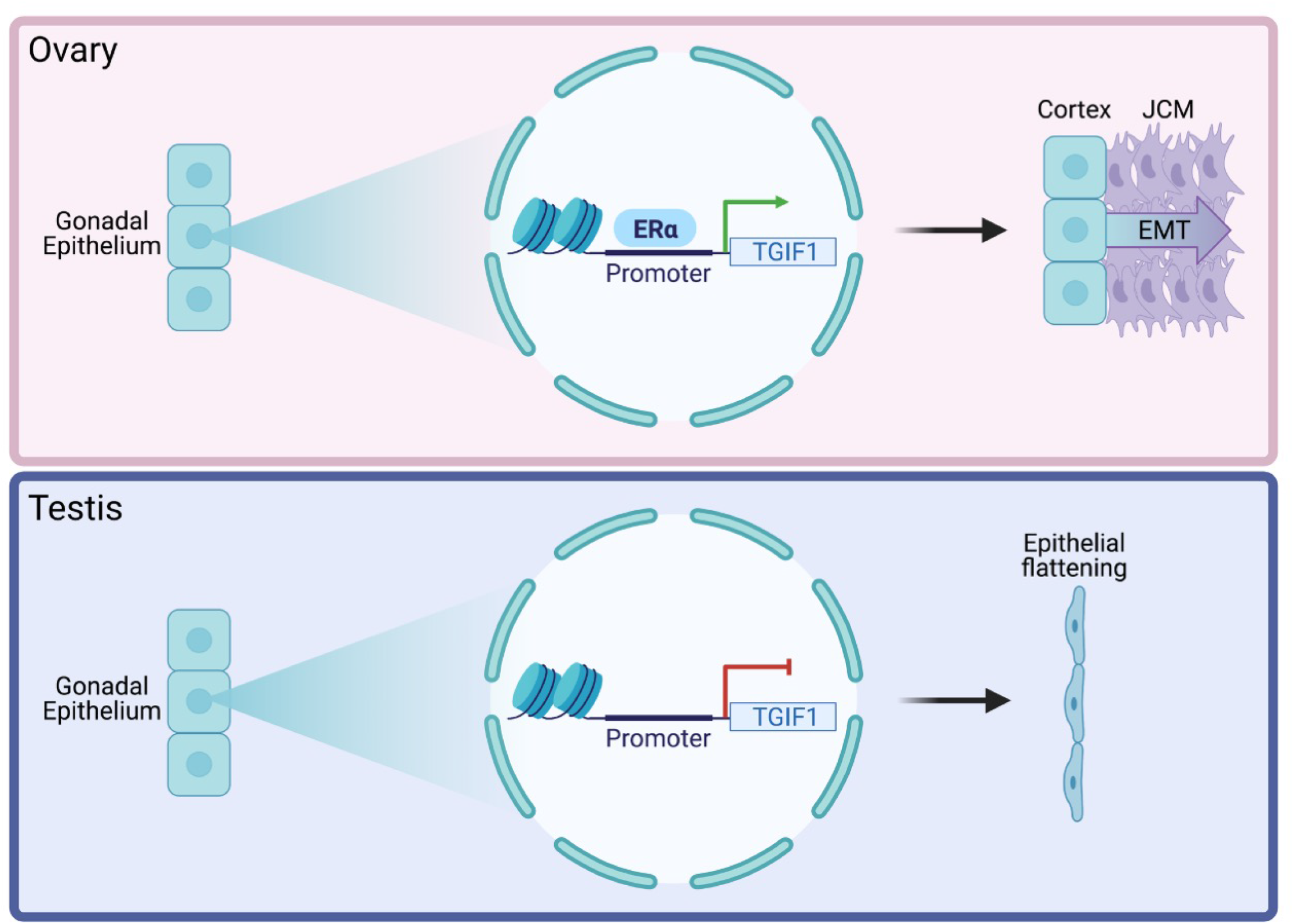
Role of TGIF1 in epithelial maintenance and juxtacortical medulla formation. In chicken ovaries (top), the activation of the ER-α signaling pathway by estrogens induces the expression of TGF1 in the gonadal epithelium, resulting in the epithelial structure maintenance and the formation of the juxtacortical medulla by increased epithelial to mesenchyme transition (EMT). In chicken testis (bottom), TGIF1 expression is not induced in the gonadal epithelial cells due to the lack of ER-α signaling. This results in the epithelial flattening and the lack juxtacortical medulla formation.

## MATERIALS AND METHODS

### Eggs and samples

Fertilized HyLine Brown chickens (*Gallus gallus domesticus*) eggs were obtained from Research Poultry Farm (Victoria, Australia) and incubated under humid conditions at 37 °C. Gonads and urogenital systems were collected at various time points throughout development and staged *in ovo* according to Hamburger and Hamilton (Hamburger and Hamilton, 1951). PCR sexing was performed as mentioned before (Clinton et al., 2001).

### qRT-PCR

Gonadal pairs were collected in Trizol reagent (Sigma-Aldrich) and kept at −80 degrees till processing. After sexing, 3 gonadal pairs from the same sex were pooled for each sample, homogenized and followed RNA extraction protocol by Phenol-Chloroform method as per the manufacturer’s instructions. Genomic DNA was removed from the RNA samples using DNA-free™ DNA Removal Kit (Invitrogen) and 200-500 ug of RNA was converted into cDNA using Promega Reverse Transcription System. qRT-PCR was performed using QuantiNova SYBR® Green PCR Kit. Expression levels were quantified by Pfaffl method (Pfaffl, 2001) using β-actin as housekeeping gene. Data was analyzed using multiple t-tests (one per embryonic stage or treatment). Statistical significance determined using the Holm-Sidak. All used primers are listed in Sup. table 1.

### Whole mount *in situ* hybridization

Whole mount in situ hybridization (WISH) was performed as previously described (Estermann et al., 2020). At least three embryos were used for each stage and sex. Urogenital systems were dissected and fixed overnight in 4% paraformaldehyde. Tissues were dehydrated in a methanol series and stored till usage. Samples then were rehydrated to PBS plus 0.1% Triton X-100 before digestion in proteinase K (1 ug/mL in PBS plus 0.1% Triton X-100) for 30 to 90 minutes at RT. Tissues were then washed, briefly refixed, and incubated overnight at 65°C in (pre)hybridization buffer. Digoxigenin-labeled antisense RNA probes were synthesized using a digoxigenin labeling kit, according to the manufacturer’s instructions (Life Technologies). TGIF1 probes (514 bp) were cloned from gonadal cDNA using primers listed in Table S1. DNA sequences were cloned into pGEM-T Easy vector and sequences were confirmed before use. For probe generation, a DNA template was first generated by PCR amplification of the insert, using M13 forward and reverse primers, encompassing RNA polymerase binding sites. Antisense and sense digoxigenin-labeled RNA probes were generated using the relevant T7 or SP6 RNA polymerase sites present in the amplified PCR product. Following synthesis, riboprobes were precipitated overnight at −20°C. For each probe, 7.5 uL were added to 2 mL (pre)hybridization mix and incubated overnight at 65 °C. Following low and high stringency washes, tissues were washed, preblocked, and incubated overnight at 4 °C with alkaline phosphatase–conjugated anti-digoxigenin antibodies in TBTX (1:2000; Roche). Following extensive washing in TBTX, tissues were exposed to chromogen (NBT/BCIP) for up to 3 hours. For each gene, the color reaction was stopped at the same time by rinsing in NTMT buffer, followed by washing in PBS and imaging. Tissues were then overstained cryoprotected in PBS plus 30% sucrose, snap frozen in OCT embedding compound, and cryosectioned between 14 and 18 um or 10 um if they were processed for immunofluorescence.

### Sex reversal

For masculinization, eggs were either injected with 1.0 mg of fadrozole (Novartis) in 100 uL of phosphate-buffered saline (PBS) or injected with PBS alone at E3.5 as previously described (Hirst et al., 2017a). For feminization 17β-estradiol (Sigma-Aldrich) was initially resuspended in 100% Ethanol (10 mg/ul) and then diluted to 1 mg/ml in sesame oil. 100 ul of this 1 mg/ml solution (0.1 mg of E2) or a 10% Ethanol in Sesame oil solution (Vehicle) was injected into E3.5 eggs. Eggs were incubated until day 9.5 of development (HH34) before processing them for whole mount *in situ* hybridization.

### TGIF1 overexpression construct design and electroporation

The Tol2 system was used to integrate TGIF1 overexpression construct into the genome of electroporated cells in the chicken embryos (Kawakami, 2007; Sato et al., 2007). TGIF1 ORF was amplified from gonadal cDNA using specific primers (See Table S1) and TA cloned into pT2-aCAGS-GcT or RCAS(A)-aCAGS-GcT and sequenced. DF1 cell (ATCC) transfection with TOL2-GFP-T2A-TGIF1 overexpression plasmid or control plasmid and transposase expressing plasmid was performed following the Lipofectamine 2000 protocol (Life Technologies). Cells were collected 48 hours post transfection and Trizol RNA extraction was performed as described before.

*In ovo* electroporation of p-CAGGS-Transposase with pT2-aCAGS-GcT-T2A-TGIF1 (TGIF1 OE) or pT2-aCAGS-GcT (GFP Control) constructs was performed as previously described (Hirst et al., 2017b) on E2.5 embryos, targeting the left coelomic epithelium. Embryos were harvested at E7.5-E8.5, sexed and processed for immunofluorescence. For RCAS electroporation, RCAS(A)-aCAGS-GcT-T2A-TGIF1 (TGIF1 OE) or RCAS(A)-aCAGS-GcT (GFP control) were electroporated.

### TGIF1 shRNA design and electroporation

TGIF1 shRNA design, validation and cloning was performed as previously described (Roly et al., 2020, Major et al., 2019). Four different shRNAs were designed against TGIF1 ORF, ranked for effectiveness (Clarke et al., 2017) and cloned into RCAS(D)-nBFP plasmid. A PCR-based amplification of the shRNA template along with the chicken U6-4 promoter was used (Lambeth et al., 2015) (Table S1). Their ability to knock down TGIF1 expression was assessed in vitro in chicken fibroblastic DF-1 cells. Firstly, DF-1 cells were transfected with the plasmids containing BFP-T2A and a non-specific shRNA (firefly sh774) (Roly et al., 2020) or with 4 different putative shRNA designed for TGIF1 knockdown (sh318, sh364, sh416 and sh998), following the Lipofectamine 2000 protocol (Life Technologies). After all cells were BFP positive, they were transfected with the TOL2-GFP-T2A-TGIF1 overexpression plasmid, a transposase expressing plasmid and a TOL2 plasmid expressing mCherry (as a transfection control) following the Lipofectamine 2000 protocol (Life Technologies). 48 hours post transfection, cells were fixed in 4% PFA for 15 minutes, stained with DAPI and imaged using a Leica AF600LX microscope. GFP-T2A-TGIF1 intensity was determined on a per cell basis using an established image analysis pipeline (Major et al., 2017). DAPI was used to identify the cell nuclei and mCherry positive cells were gated for further analysis. TGIF1 sh998 showed the stronger TGIF1/GFP inhibition and was cloned into a TOL2 vector expressing nuclear BFP. TOL2-TGIF1sh998-nBFP or TOL2-Fireflysh774-nBFP (non-silencing shRNA) was *in ovo* co-electroporated with a plasmid expressing transposase and TOL2-ACAGS-GFP (electroporation reporter) into the left coelomic epithelium at E2.5. Urogenital systems were collected at E8.5, sexed and immunofluorescence was performed against different gonadal markers.

### Immunofluorescence

At least three embryos per time point and/or treatment were examined. Tissues were fixed in 4% paraformaldehyde/PBS for 15 minutes at room temperature. Tissues were cryoprotected in PBS plus 30% sucrose, snap frozen in OCT embedding compound, and sectioned at 10 um. Some slides were first subjected to antigen retrieval, using an automated system, the Dako PT Link. Slides were firstly baked at 60 °C for 30 minutes. Retrieval was then performed with the Dako Target retrieval solution, a citrate-based (pH 6.0). Slides were then placed in the retrieval machine and retrieved at 98 °C for 30 minutes. All sections were permeabilized in PBS containing 1% Triton X-100, blocked in PBS 2% BSA for 1 hour, and incubated ON at 4 °C with primary antibodies in 1% BSA in PBS. Primary antibodies used: goat anti-GFP (Rockland 600-101-215, 1:500), mouse anti-pan-cytokeratin (Novus Bio NBP2-29429, 1:200), rabbit anti-DMRT1 (in house antibody, 1:2000), rabbit anti-SOX9 (Millipore antibody AB5535, 1:4000), rabbit anti-AMH (Abexa ABX132175, 1:1000), rabbit anti-aromatase (in house antibody, 1:5000), mouse anti-fibronectin (Serotec 4470–4339, 1:500) and rabbit anti-CVH (in house antibody 1:500). Alexa Fluor secondary antibodies were used (donkey or goat anti-rabbit, mouse or goat 488 or 594; Life technologies). Sections were counterstained with DAPI and mounted in Fluorsave (Milipore). For WISH samples, sections were processed for antigen retrieval (as mentioned above). After the secondary antibody incubation, sections were treated with 0.3% Sudan Black (w/v) in 70% EtOH (v/v) for 10 minutes followed by 8 quick PBS washes. Sections were counterstained with DAPI and mounted.

### Image quantification

Gonadal, epithelial, medulla and juxtacortical medulla area were manually quantified using Fiji (Schindelin et al., 2012). Epithelial average thickness was calculated by dividing the epithelial area over the length of the epithelium.

## ACKNOWLEDGEMENTS

The authors acknowledge use of the facilities and technical assistance of Monash Histology Platform, Department of Anatomy and Developmental Biology, Monash University. The authors acknowledge the facilities and technical assistance of Monash Micro Imaging.

## REFERENCES

Andrews, J. E., Smith, C. A. & Sinclair, A. H. 1997. Sites of estrogen receptor and aromatase expression in the chicken embryo. Gen Comp Endocrinol, 108, 182–90.

Ayers, K. L., Lambeth, L. S., Davidson, N. M., Sinclair, A. H., Oshlack, A. & Smith, C. A. 2015. Identification of candidate gonadal sex differentiation genes in the chicken embryo using RNA-seq. BMC Genomics, 16, 704.

Ayers, K. L., Sinclair, A. H. & Smith, C. A. 2013. The molecular genetics of ovarian differentiation in the avian model. Sex Dev, 7, 80–94.

Ayyar, S., Jiang, J., Collu, A., White-Cooper, H. & White, R. A. 2003. Drosophila TGIF is essential for developmentally regulated transcription in spermatogenesis. Development, 130, 2841-52.

Bannister, S. C., Smith, C. A., Roeszler, K. N., Doran, T. J., Sinclair, A. H. & Tizard, M. L. 2011. Manipulation of estrogen synthesis alters MIR202* expression in embryonic chicken gonads. Biol Reprod, 85, 22–30.

Barrios, F., Filipponi, D., Pellegrini, M., Paronetto, M. P., Di Siena, S., Geremia, R., Rossi, P., De Felici, M., Jannini, E. A. & Dolci, S. 2010. Opposing effects of retinoic acid and FGF9 on Nanos2 expression and meiotic entry of mouse germ cells. J Cell Sci, 123, 871–80.

Bertolino, E., Reimund, B., Wildt-Perinic, D. & Clerc, R. G. 1995. A novel homeobox protein which recognizes a TGT core and functionally interferes with a retinoid-responsive motif. J Biol Chem, 270, 31178–88.

Bowles, J., Feng, C. W., Spiller, C., Davidson, T. L., Jackson, A. & Koopman, P. 2010. FGF9 suppresses meiosis and promotes male germ cell fate in mice. Dev Cell, 19, 440–9.

Brennan, J. & Capel, B. 2004. One tissue, two fates: molecular genetic events that underlie testis versus ovary development. Nat Rev Genet, 5, 509–21.

Capel, B. 2017. Vertebrate sex determination: evolutionary plasticity of a fundamental switch. Nat Rev Genet, 18, 675–689.

Castillo, H. A., Cravo, R. M., Azambuja, A. P., Simoes-Costa, M. S., Sura-Trueba, S., Gonzalez, J., Slonimsky, E., Almeida, K., Abreu, J. G., De Almeida, M. A., Sobreira, T. P., De Oliveira, S. H., De Oliveira, P. S., Signore, I. A., Colombo, A., Concha, M. L., Spengler, T. S., Bronner-Fraser, M., Nobrega, M., Rosenthal, N. & Xavier-Neto, J. 2010. Insights into the organization of dorsal spinal cord pathways from an evolutionarily conserved raldh2 intronic enhancer. Development, 137, 507–18.

Chassot, A. A., Ranc, F., Gregoire, E. P., Roepers-Gajadien, H. L., Taketo, M. M., Camerino, G., De Rooij, D. G., Schedl, A. & Chaboissier, M. C. 2008. Activation of beta-catenin signaling by Rspo1 controls differentiation of the mammalian ovary. Hum Mol Genet, 17, 1264–77.

Chen, M., Zhang, L., Cui, X., Lin, X., Li, Y., Wang, Y., Wang, Y., Qin, Y., Chen, D., Han, C., Zhou, B., Huff, V. & Gao, F. 2017. Wt1 directs the lineage specification of sertoli and granulosa cells by repressing Sf1 expression. Development, 144, 44–53.

Clarke, B. D., Mccoll, K. A., Ward, A. C. & Doran, T. J. 2017. shRNAs targeting either the glycoprotein or polymerase genes inhibit Viral haemorrhagic septicaemia virus replication in zebrafish ZF4 cells. Antiviral Res, 141, 124–132.

Clinton, M., Haines, L., Belloir, B. & Mcbride, D. 2001. Sexing chick embryos: a rapid and simple protocol. Br Poult Sci, 42, 134–8.

Colvin, J. S., Green, R. P., Schmahl, J., Capel, B. & Ornitz, D. M. 2001. Male-to-female sex reversal in mice lacking fibroblast growth factor 9. Cell, 104, 875–89.

Defalco, T., Takahashi, S. & Capel, B. 2011. Two distinct origins for Leydig cell progenitors in the fetal testis. Dev Biol, 352, 14–26.

Dinapoli, L., Batchvarov, J. & Capel, B. 2006. FGF9 promotes survival of germ cells in the fetal testis. Development, 133, 1519–27.

Elbrecht, A. & Smith, R. G. 1992. Aromatase enzyme activity and sex determination in chickens. Science, 255, 467–70.

Estermann, M. A., Williams, S., Hirst, C. E., Roly, Z. Y., Serralbo, O., Adhikari, D., Powell, D., Major, A. T. & Smith, C. A. 2020. Insights into Gonadal Sex Differentiation Provided by Single-Cell Transcriptomics in the Chicken Embryo. Cell Rep, 31, 107491.

Fujimoto, Y., Tanaka, S. S., Yamaguchi, Y. L., Kobayashi, H., Kuroki, S., Tachibana, M., Shinomura, M., Kanai, Y., Morohashi, K., Kawakami, K. & Nishinakamura, R. 2013. Homeoproteins Six1 and Six4 regulate male sex determination and mouse gonadal development. Dev Cell, 26, 416–30.

Gandhi, S., Ezin, M. & Bronner, M. E. 2020. Reprogramming Axial Level Identity to Rescue Neural-Crest-Related Congenital Heart Defects. Dev Cell, 53, 300–315 e4.

Gasc, J. M. & Stumpf, W. E. 1981. Sexual differentiation of the urogenital tract in the chicken embryo: autoradiographic localization of sex-steroid target cells during development. J Embryol Exp Morphol, 63, 207–23.

Gonen, N. & Lovell-Badge, R. 2019. The regulation of Sox9 expression in the gonad. Curr Top Dev Biol, 134, 223–252.

Gonen, N., Quinn, A., O’neill, H. C., Koopman, P. & Lovell-Badge, R. 2017. Normal Levels of Sox9 Expression in the Developing Mouse Testis Depend on the TES/TESCO Enhancer, but This Does Not Act Alone. PLoS Genet, 13, e1006520.

Gongal, P. A. & Waskiewicz, A. J. 2008. Zebrafish model of holoprosencephaly demonstrates a key role for TGIF in regulating retinoic acid metabolism. Hum Mol Genet, 17, 525–38.

Gonzalez-Moran, M. G. 2014. Changes in the cellular localization of estrogen receptor alpha in the growing and regressing ovaries of Gallus domesticus during development. Biochem Biophys Res Commun, 447, 197–204.

Guioli, S. & Lovell-Badge, R. 2007. PITX2 controls asymmetric gonadal development in both sexes of the chick and can rescue the degeneration of the right ovary. Development, 134, 4199–208.

Guioli, S., Nandi, S., Zhao, D., Burgess-Shannon, J., Lovell-Badge, R. & Clinton, M. 2014. Gonadal asymmetry and sex determination in birds. Sex Dev, 8, 227–42.

Guioli, S., Zhao, D., Nandi, S., Clinton, M. & Lovell-Badge, R. 2020. Oestrogen in the chick embryo can induce chromosomally male ZZ left gonad epithelial cells to form an ovarian cortex that can support oogenesis. Development, 147.

Gustin, S. E., Stringer, J. M., Hogg, K., Sinclair, A. H. & Western, P. S. 2016. FGF9, activin and TGFbeta promote testicular characteristics in an XX gonad organ culture model. Reproduction, 152, 529–43.

Hacker, A., Capel, B., Goodfellow, P. & Lovell-Badge, R. 1995. Expression of Sry, the mouse sex determining gene. Development, 121, 1603–14.

Haider, M. T., Saito, H., Zarrer, J., Uzhunnumpuram, K., Nagarajan, S., Kari, V., Horn-Glander, M., Werner, S., Hesse, E. & Taipaleenmaki, H. 2020. Breast cancer bone metastases are attenuated in a Tgif1-deficient bone microenvironment. Breast Cancer Res, 22, 34.

Hamburger, V. & Hamilton, H. L. 1951. A series of normal stages in the development of the chick embryo. J Morphol, 88, 49–92.

Hatano, A., Matsumoto, M., Higashinakagawa, T. & Nakayama, K. I. 2010. Phosphorylation of the chromodomain changes the binding specificity of Cbx2 for methylated histone H3. Biochem Biophys Res Commun, 397, 93–9.

Hirst, C. E., Major, A. T., Ayers, K. L., Brown, R. J., Mariette, M., Sackton, T. B. & Smith, C. A. 2017a. Sex Reversal and Comparative Data Undermine the W Chromosome and Support Z-linked DMRT1 as the Regulator of Gonadal Sex Differentiation in Birds. Endocrinology, 158, 2970–2987.

Hirst, C. E., Serralbo, O., Ayers, K. L., Roeszler, K. N. & Smith, C. A. 2017b. Genetic Manipulation of the Avian Urogenital System Using In Ovo Electroporation. In: Sheng, G. (ed.) Avian and Reptilian Developmental Biology: Methods and Protocols. New York, NY: Springer New York.

Hu, S., Zhu, M., Wang, J., Li, L., He, H., Hu, B., Hu, J. & Xia, L. 2021. Histomorphology and gene expression profiles during early ovarian folliculogenesis in duck and goose. Poult Sci, 100, 1098–1108.

Hu, Y., Yu, H., Shaw, G., Renfree, M. B. & Pask, A. J. 2011. Differential roles of TGIF family genes in mammalian reproduction. BMC Dev Biol, 11, 58.

Ioannidis, J., Taylor, G., Zhao, D., Liu, L., Idoko-Akoh, A., Gong, D., Lovell-Badge, R., Guioli, S., Mcgrew, M. & Clinton, M. 2020. Primary sex determination in chickens depends on DMRT1 dosage, but gonadal sex does not determine secondary sexual characteristics in adult birds. bioRxiv, 2020.09.18.303040.

Ioannidis, J., Taylor, G., Zhao, D., Liu, L., Idoko-Akoh, A., Gong, D., Lovell-Badge, R., Guioli, S., Mcgrew, M. J. & Clinton, M. 2021. Primary sex determination in birds depends on DMRT1 dosage, but gonadal sex does not determine adult secondary sex characteristics. Proc Natl Acad Sci U S A, 118.

Ishimaru, Y., Komatsu, T., Kasahara, M., Katoh-Fukui, Y., Ogawa, H., Toyama, Y., Maekawa, M., Toshimori, K., Chandraratna, R. A., Morohashi, K. & Yoshioka, H. 2008. Mechanism of asymmetric ovarian development in chick embryos. Development, 135, 677–85.

Johnson, A. L. & Woods, D. C. 2009. Dynamics of avian ovarian follicle development: cellular mechanisms of granulosa cell differentiation. Gen Comp Endocrinol, 163, 12–7.

Jordan, B. K., Shen, J. H., Olaso, R., Ingraham, H. A. & Vilain, E. 2003. Wnt4 overexpression disrupts normal testicular vasculature and inhibits testosterone synthesis by repressing steroidogenic factor 1/beta-catenin synergy. Proc Natl Acad Sci U S A, 100, 10866–71.

Karl, J. & Capel, B. 1998. Sertoli cells of the mouse testis originate from the coelomic epithelium. Dev Biol, 203, 323–33.

Kashimada, K. & Koopman, P. 2010. Sry: the master switch in mammalian sex determination. Development, 137, 3921–30.

Kashimada, K., Pelosi, E., Chen, H., Schlessinger, D., Wilhelm, D. & Koopman, P. 2011. FOXL2 and BMP2 act cooperatively to regulate follistatin gene expression during ovarian development. Endocrinology, 152, 272–80.

Kent, J., Wheatley, S. C., Andrews, J. E., Sinclair, A. H. & Koopman, P. 1996. A male-specific role for SOX9 in vertebrate sex determination. Development, 122, 2813–22.

Kim, Y., Bingham, N., Sekido, R., Parker, K. L., Lovell-Badge, R. & Capel, B. 2007. Fibroblast growth factor receptor 2 regulates proliferation and Sertoli differentiation during male sex determination. Proc Natl Acad Sci U S A, 104, 16558–63.

Kim, Y., Kobayashi, A., Sekido, R., Dinapoli, L., Brennan, J., Chaboissier, M. C., Poulat, F., Behringer, R. R., Lovell-Badge, R. & Capel, B. 2006. Fgf9 and Wnt4 act as antagonistic signals to regulate mammalian sex determination. PLoS Biol, 4, e187.

Koopman, P., Gubbay, J., Vivian, N., Goodfellow, P. & Lovell-Badge, R. 1991. Male development of chromosomally female mice transgenic for Sry. Nature, 351, 117–21.

Kusaka, M., Katoh-Fukui, Y., Ogawa, H., Miyabayashi, K., Baba, T., Shima, Y., Sugiyama, N., Sugimoto, Y., Okuno, Y., Kodama, R., Iizuka-Kogo, A., Senda, T., Sasaoka, T., Kitamura, K., Aizawa, S. & Morohashi, K. 2010. Abnormal epithelial cell polarity and ectopic epidermal growth factor receptor (EGFR) expression induced in Emx2 KO embryonic gonads. Endocrinology, 151, 5893–904.

Lambeth, L. S., Ayers, K., Cutting, A. D., Doran, T. J., Sinclair, A. H. & Smith, C. A. 2015. Anti-Mullerian Hormone Is Required for Chicken Embryonic Urogenital System Growth but Not Sexual Differentiation. Biol Reprod, 93, 138.

Lambeth, L. S., Raymond, C. S., Roeszler, K. N., Kuroiwa, A., Nakata, T., Zarkower, D. & Smith, C. A. 2014. Over-expression of DMRT1 induces the male pathway in embryonic chicken gonads. Dev Biol, 389, 160–72.

Lau, Y. F. & Li, Y. 2009. The human and mouse sex-determining SRY genes repress the Rspol/beta-catenin signaling. J Genet Genomics, 36, 193–202.

Lavery, R., Chassot, A. A., Pauper, E., Gregoire, E. P., Klopfenstein, M., De Rooij, D. G., Mark, M., Schedl, A., Ghyselinck, N. B. & Chaboissier, M. C. 2012. Testicular differentiation occurs in absence of R-spondin1 and Sox9 in mouse sex reversals. PLoS Genet, 8, e1003170.

Lee, B. K., Shen, W., Lee, J., Rhee, C., Chung, H., Kim, K. Y., Park, I. H. & Kim, J. 2015. Tgif1 Counterbalances the Activity of Core Pluripotency Factors in Mouse Embryonic Stem Cells. Cell Rep, 13, 52–60.

Li, J., Zhao, D., Guo, C., Li, J., Mi, Y. & Zhang, C. 2016. Involvement of Notch signaling in early chick ovarian follicle development. Cell Biol Int, 40, 65–73.

Li, Y., Zhang, L., Hu, Y., Chen, M., Han, F., Qin, Y., Chen, M., Cui, X., Duo, S., Tang, F. & Gao, F. 2017. beta-Catenin directs the transformation of testis Sertoli cells to ovarian granulosa-like cells by inducing Foxl2 expression. J Biol Chem, 292, 17577–17586.

Li, Y., Zheng, M. & Lau, Y. F. 2014. The sex-determining factors SRY and SOX9 regulate similar target genes and promote testis cord formation during testicular differentiation. Cell Rep, 8, 723–33.

Lin, Y. T., Barske, L., Defalco, T. & Capel, B. 2017. Numb regulates somatic cell lineage commitment during early gonadogenesis in mice. Development, 144, 1607–1618.

Lin, Y. T. & Capel, B. 2015. Cell fate commitment during mammalian sex determination. Curr Opin Genet Dev, 32, 144–52.

Liu, C., Peng, J., Matzuk, M. M. & Yao, H. H. 2015. Lineage specification of ovarian theca cells requires multicellular interactions via oocyte and granulosa cells. Nat Commun, 6, 6934.

Liu, X., Zhang, H., Gao, L., Yin, Y., Pan, X., Li, Z., Li, N., Li, H. & Yu, Z. 2014. Negative interplay of retinoic acid and TGF-beta signaling mediated by TG-interacting factor to modulate mouse embryonic palate mesenchymal-cell proliferation. Birth Defects Res B Dev Reprod Toxicol, 101, 403–9.

Lorda-Diez, C. I., Montero, J. A., Martinez-Cue, C., Garcia-Porrero, J. A. & Hurle, J. M. 2009. Transforming growth factors beta coordinate cartilage and tendon differentiation in the developing limb mesenchyme. J Biol Chem, 284, 29988–96.

Maatouk, D. M., Dinapoli, L., Alvers, A., Parker, K. L., Taketo, M. M. & Capel, B. 2008. Stabilization of beta-catenin in XY gonads causes male-to-female sex-reversal. Hum Mol Genet, 17, 2949–55.

Major, A. T., Ayers, K., Chue, J., Roeszler, K. & Smith, C. 2019. FOXL2 antagonises the male developmental pathway in embryonic chicken gonads. J Endocrinol.

Major, A. T., Miyamoto, Y., Lo, C. Y., Jans, D. A. & Loveland, K. L. 2017. Development of a pipeline for automated, high-throughput analysis of paraspeckle proteins reveals specific roles for importin alpha proteins. Sci Rep, 7, 43323.

Melhuish, T. A. & Wotton, D. 2000. The interaction of the carboxyl terminus-binding protein with the Smad corepressor TGIF is disrupted by a holoprosencephaly mutation in TGIF. J Biol Chem, 275, 39762–6.

Mendez, C., Alcantara, L., Escalona, R., Lopez-Casillas, F. & Pedernera, E. 2006. Transforming growth factor beta inhibits proliferation of somatic cells without influencing germ cell number in the chicken embryonic ovary. Cell Tissue Res, 325, 143–9.

Munger, S. C. & Capel, B. 2012. Sex and the circuitry: progress toward a systems-level understanding of vertebrate sex determination. Wiley Interdiscip Rev Syst Biol Med, 4, 401–12.

Nef, S., Stevant, I. & Greenfield, A. 2019. Characterizing the bipotential mammalian gonad. Curr Top Dev Biol, 134, 167–194.

Nicol, B. & Yao, H. H. 2014. Building an ovary: insights into establishment of somatic cell lineages in the mouse. Sex Dev, 8, 243–51.

Niu, W. & Spradling, A. C. 2020. Two distinct pathways of pregranulosa cell differentiation support follicle formation in the mouse ovary. Proc Natl Acad Sci U S A, 117, 20015–20026.

Omotehara, T., Minami, K., Mantani, Y., Umemura, Y., Nishida, M., Hirano, T., Yoshioka, H., Kitagawa, H., Yokoyama, T. & Hoshi, N. 2017. Contribution of the coelomic epithelial cells specific to the left testis in the chicken embryo. Dev Dyn, 246, 148–156.

Parma, P., Radi, O., Vidal, V., Chaboissier, M. C., Dellambra, E., Valentini, S., Guerra, L., Schedl, A. & Camerino, G. 2006. R-spondin1 is essential in sex determination, skin differentiation and malignancy. Nat Genet, 38, 1304–9.

Pfaffl, M. W. 2001. A new mathematical model for relative quantification in real-time RT-PCR. Nucleic Acids Res, 29, e45.

Pieau, C. & Dorizzi, M. 2004. Oestrogens and temperature-dependent sex determination in reptiles: all is in the gonads. J Endocrinol, 181, 367–77.

Powers, S. E., Taniguchi, K., Yen, W., Melhuish, T. A., Shen, J., Walsh, C. A., Sutherland, A. E. & Wotton, D. 2010. Tgif1 and Tgif2 regulate Nodal signaling and are required for gastrulation. Development, 137, 249–59.

Qin, Y. & Bishop, C. E. 2005. Sox9 is sufficient for functional testis development producing fertile male mice in the absence of Sry. Hum Mol Genet, 14, 1221–9.

Rodriguez-Leon, J., Rodriguez Esteban, C., Marti, M., Santiago-Josefat, B., Dubova, I., Rubiralta, X. & Izpisua Belmonte, J. C. 2008. Pitx2 regulates gonad morphogenesis. Proc Natl Acad Sci U S A, 105, 11242–7.

Roly, Z. Y., Major, A. T., Fulcher, A., Estermann, M. A., Hirst, C. E. & Smith, C. A. 2020. Adhesion G-protein-coupled receptor, GPR56, is required for Mullerian duct development in the chick. J Endocrinol, 244, 395–413.

Rotgers, E., Jorgensen, A. & Yao, H. H. 2018. At the Crossroads of Fate-Somatic Cell Lineage Specification in the Fetal Gonad. Endocr Rev, 39, 739–759.

Sato, Y., Kasai, T., Nakagawa, S., Tanabe, K., Watanabe, T., Kawakami, K. & Takahashi, Y. 2007. Stable integration and conditional expression of electroporated transgenes in chicken embryos. Dev Biol, 305, 616–24.

Scheib, D. 1983. Effects and role of estrogens in avian gonadal differentiation. Differentiation. 1983/01/01 ed.

Schindelin, J., Arganda-Carreras, I., Frise, E., Kaynig, V., Longair, M., Pietzsch, T., Preibisch, S., Rueden, C., Saalfeld, S., Schmid, B., Tinevez, J. Y., White, D. J., Hartenstein, V., Eliceiri, K., Tomancak, P. & Cardona, A. 2012. Fiji: an open-source platform for biological-image analysis. Nat Methods, 9, 676–82.

Schmahl, J., Kim, Y., Colvin, J. S., Ornitz, D. M. & Capel, B. 2004. Fgf9 induces proliferation and nuclear localization of FGFR2 in Sertoli precursors during male sex determination. Development, 131, 3627–36.

Schmid, M., Smith, J., Burt, D. W., Aken, B. L., Antin, P. B., Archibald, A. L., Ashwell, C., Blackshear, P. J., Boschiero, C., Brown, C. T., Burgess, S. C., Cheng, H. H., Chow, W., Coble, D. J., Cooksey, A., Crooijmans, R. P., Damas, J., Davis, R. V., De Koning, D. J., Delany, M. E., Derrien, T., Desta, T. T., Dunn, I. C., Dunn, M., Ellegren, H., Eory, L., Erb, I., Farre, M., Fasold, M., Fleming, D., Flicek, P., Fowler, K. E., Fresard, L., Froman, D. P., Garceau, V., Gardner, P. P., Gheyas, A. A., Griffin, D. K., Groenen, M. A., Haaf, T., Hanotte, O., Hart, A., Hasler, J., Hedges, S. B., Hertel, J., Howe, K., Hubbard, A., Hume, D. A., Kaiser, P., Kedra, D., Kemp, S. J., Klopp, C., Kniel, K. E., Kuo, R., Lagarrigue, S., Lamont, S. J., Larkin, D. M., Lawal, R. A., Markland, S. M., Mccarthy, F., Mccormack, H. A., Mcpherson, M. C., Motegi, A., Muljo, S. A., Munsterberg, A., Nag, R., Nanda, I., Neuberger, M., Nitsche, A., Notredame, C., Noyes, H., O’Connor, R., O’Hare, E. A., Oler, A. J., Ommeh, S. C., Pais, H., Persia, M., Pitel, F., Preeyanon, L., Prieto Barja, P., Pritchett, E. M., Rhoads, D. D., Robinson, C. M., Romanov, M. N., Rothschild, M., Roux, P. F., Schmidt, C. J., Schneider, A. S., Schwartz, M. G., Searle, S. M., Skinner, M. A., Smith, C. A., Stadler, P. F., Steeves, T. E., Steinlein, C., Sun, L., Takata, M., Ulitsky, I., Wang, Q., Wang, Y., et al. 2015. Third Report on Chicken Genes and Chromosomes 2015. Cytogenet Genome Res, 145, 78–179.

Sekido, R., Bar, I., Narvaez, V., Penny, G. & Lovell-Badge, R. 2004. SOX9 is up-regulated by the transient expression of SRY specifically in Sertoli cell precursors. Dev Biol, 274, 271–9.

Sekido, R. & Lovell-Badge, R. 2007. Mechanisms of gonadal morphogenesis are not conserved between chick and mouse. Dev Biol, 302, 132–42.

Sekido, R. & Lovell-Badge, R. 2008. Sex determination involves synergistic action of SRY and SF1 on a specific Sox9 enhancer. Nature, 453, 930–4.

Shen, J. & Walsh, C. A. 2005. Targeted disruption of Tgif, the mouse ortholog of a human holoprosencephaly gene, does not result in holoprosencephaly in mice. Mol Cell Biol, 25, 3639–47.

Sinclair, A. H., Berta, P., Palmer, M. S., Hawkins, J. R., Griffiths, B. L., Smith, M. J., Foster, J. W., Frischauf, A. M., Lovell-Badge, R. & Goodfellow, P. N. 1990. A gene from the human sex-determining region encodes a protein with homology to a conserved DNA-binding motif. Nature, 346, 240–4.

Smith, C. A., Katz, M. & Sinclair, A. H. 2003. DMRT1 is upregulated in the gonads during female-to-male sex reversal in ZW chicken embryos. Biol Reprod, 68, 560–70.

Smith, C. A., Roeszler, K. N., Bowles, J., Koopman, P. & Sinclair, A. H. 2008. Onset of meiosis in the chicken embryo; evidence of a role for retinoic acid. BMC Dev Biol, 8, 85.

Smith, C. A., Roeszler, K. N., Ohnesorg, T., Cummins, D. M., Farlie, P. G., Doran, T. J. & Sinclair, A. H. 2009. The avian Z-linked gene DMRT1 is required for male sex determination in the chicken. Nature, 461, 267–71.

Smith, C. A. & Sinclair, A. H. 2004. Sex determination: insights from the chicken. Bioessays, 26, 120–32.

Spiller, C., Koopman, P. & Bowles, J. 2017. Sex Determination in the Mammalian Germline. Annu Rev Genet, 51, 265–285.

Stevant, I., Kuhne, F., Greenfield, A., Chaboissier, M. C., Dermitzakis, E. T. & Nef, S. 2019. Dissecting Cell Lineage Specification and Sex Fate Determination in Gonadal Somatic Cells Using Single-Cell Transcriptomics. Cell Rep, 26, 3272–3283 e3.

Stevant, I. & Nef, S. 2019. Genetic Control of Gonadal Sex Determination and Development. Trends Genet, 35, 346–358.

Stevant, I., Neirijnck, Y., Borel, C., Escoffier, J., Smith, L. B., Antonarakis, S. E., Dermitzakis, E. T. & Nef, S. 2018. Deciphering Cell Lineage Specification during Male Sex Determination with Single-Cell RNA Sequencing. Cell Rep, 22, 1589–1599.

Sun, W., Cai, H., Zhang, G., Zhang, H., Bao, H., Wang, L., Ye, J., Qian, G. & Ge, C. 2017. Dmrt1 is required for primary male sexual differentiation in Chinese soft-shelled turtle Pelodiscus sinensis. Sci Rep, 7, 4433.

Svingen, T. & Koopman, P. 2013. Building the mammalian testis: origins, differentiation, and assembly of the component cell populations. Genes Dev, 27, 2409–26.

Tomizuka, K., Horikoshi, K., Kitada, R., Sugawara, Y., Iba, Y., Kojima, A., Yoshitome, A., Yamawaki, K., Amagai, M., Inoue, A., Oshima, T. & Kakitani, M. 2008. R-spondin1 plays an essential role in ovarian development through positively regulating Wnt-4 signaling. Hum Mol Genet, 17, 1278–91.

Ukeshima, A. 1996. Germ cell death in the degenerating right ovary of the chick embryo. Zoolog Sci, 13, 559–63.

Vaillant, S., Dorizzi, M., Pieau, C. & Richard-Mercier, N. 2001a. Sex reversal and aromatase in chicken. J Exp Zool, 290, 727–40.

Vaillant, S., Magre, S., Dorizzi, M., Pieau, C. & Richard-Mercier, N. 2001b. Expression of AMH, SF1, and SOX9 in gonads of genetic female chickens during sex reversal induced by an aromatase inhibitor. Dev Dyn, 222, 228–37.

Vidal, V. P., Chaboissier, M. C., De Rooij, D. G. & Schedl, A. 2001. Sox9 induces testis development in XX transgenic mice. Nat Genet, 28, 216–7.

Wang, J. L., Qi, Z., Li, Y. H., Zhao, H. M., Chen, Y. G. & Fu, W. 2017. TGFbeta induced factor homeobox 1 promotes colorectal cancer development through activating Wnt/beta-catenin signaling. Oncotarget, 8, 70214–70225.

Wartenberg, H., Lenz, E. & Schweikert, H. U. 1992. Sexual differentiation and the germ cell in sex reversed gonads after aromatase inhibition in the chicken embryo. Andrologia, 24, 1–6.

Wear, H. M., Eriksson, A., Yao, H. H. & Watanabe, K. H. 2017. Cell-based computational model of early ovarian development in mice. Biol Reprod, 97, 365–377.

Wotton, D., Knoepfler, P. S., Laherty, C. D., Eisenman, R. N. & Massague, J. 2001. The Smad transcriptional corepressor TGIF recruits mSin3. Cell Growth Differ, 12, 457–63.

Wotton, D., Lo, R. S., Lee, S. & Massague, J. 1999a. A Smad transcriptional corepressor. Cell, 97, 29–39.

Wotton, D., Lo, R. S., Swaby, L. A. & Massague, J. 1999b. Multiple modes of repression by the Smad transcriptional corepressor TGIF. J Biol Chem, 274, 37105–10.

Wu, Q., Kanata, K., Saba, R., Deng, C. X., Hamada, H. & Saga, Y. 2013. Nodal/activin signalingpromotes male germ cell fate and suppresses female programming in somatic cells. Development, 140, 291–300.

Xiang, G., Yi, Y., Weiwei, H. & Weiming, W. 2015. TGIF1 promoted the growth and migration of cancer cells in nonsmall cell lung cancer. Tumour Biol, 36, 9303–10.

Yao, H. H., Whoriskey, W. & Capel, B. 2002. Desert Hedgehog/Patched 1 signaling specifies fetal Leydig cell fate in testis organogenesis. Genes Dev, 16, 1433–40.

Zhang, L., Chen, M., Wen, Q., Li, Y., Wang, Y., Wang, Y., Qin, Y., Cui, X., Yang, L., Huff, V. & Gao, F. 2015a. Reprogramming of Sertoli cells to fetal-like Leydig cells by Wt1 ablation. Proc Natl Acad Sci U S A, 112, 4003–8.

Zhang, M. Z., Ferrigno, O., Wang, Z., Ohnishi, M., Prunier, C., Levy, L., Razzaque, M., Horne, W. C., Romero, D., Tzivion, G., Colland, F., Baron, R. & Atfi, A. 2015b. TGIF governs a feed-forward network that empowers Wnt signaling to drive mammary tumorigenesis. Cancer Cell, 27, 547–60.

